# A new cell type facilitated live-bearing and multiple independent origins of placentation in fish

**DOI:** 10.64898/2026.11.11.765921

**Authors:** Diego Safian, Sophie Kraunsoe, Marcin Falis, Leann Labra, David Reznick, Margarida Cardoso-Moreira

## Abstract

The origin of new organs has fascinated biologists since Darwin. Yet the molecular and evolutionary steps that give rise to new organs remain difficult to resolve, largely because most organs have ancient origins. The fish family Poeciliidae offers a rare exception. Within poeciliids, placentas have independently evolved at least nine times, including within the last∼10 million years. Here, we study the molecular and cellular drivers of five independent origins of a placenta, using pairs of sister species with and without a placenta. We identified a new maternal cell type that emerged in the common ancestor of poeciliids and likely enabled the family to transition from egg-laying to live-bearing through egg retention. This new cell type originated by partly co-opting the existing gene regulatory network of another ovarian cell type. Subsequently, this same new maternal cell type, together with the embryonic yolk sac, was repeatedly and independently co-opted to form a placenta in all five origins. The evolution of these placentas involved high levels of convergence at the pathway level (e.g., lipid metabolism and vesicle-mediated transport), although these functional changes relied primarily on distinct genes across species. Our analyses show how the evolution and modification of a new cell type enabled the evolution of live birth and placentation, providing the cellular and genetic foundations for the repeated evolution of a new organ.

---

The evolution of new organs, such as lungs, eyes, and stomachs, has enabled organisms to adapt to new ecological niches and diversify (1). Despite their importance to our understanding of evolution, the molecular and developmental mechanisms behind the origin of new organs remain poorly understood (2–4). A major challenge is that most organs originated deep in evolutionary time, and so the molecular and developmental changes that drove their emergence have long been obscured by hundreds of millions of years of morphological, physiological and molecular evolution. However, exceptions exist, and evolutionarily young organs offer an opportunity to reconstruct the molecular and developmental paths that led to their emergence. Among the best examples of newly evolved organs are vertebrate placentas.

A placenta is defined functionally as an organ made by the close association of maternal and embryonic tissues to enable nutrient transfer from the mother to the developing embryo (the known exception is Syngnathidae fishes, where the placenta is paternal) (4–6). Placentas have evolved independently many times across vertebrate lineages, including once in mammals, and many times in reptiles, amphibians, and fishes (6, 7).

Among vertebrates, placentas in the fish family Poeciliidae are one of the most compelling examples of recent organ evolution, with some originating only ∼10 million years ago (mya). Poeciliids are notable for their reproductive innovations. The family’s last common ancestor transitioned from egg-laying to live-bearing, which is thought to have driven the diversification of the family (8). In most poeciliids, live-bearing takes the form of egg-retention: mothers deposit the nutrients required for embryo development in the eggs before fertilization (as yolk), but unlike egg-laying species, eggs are retained individually inside the ovarian follicles, where they are fertilized and where embryos develop (9–11). Egg retention is an evolutionary innovation that required major adaptations of the ovarian follicle, including the loss of ovulation. The nature of these adaptations and how they arose remain unknown. Among poeciliids, some species innovated further, resulting in at least nine independent transitions from egg retention to placentation (12–14). In placental species, most maternal provisioning occurs after fertilization, with nutrients delivered directly to developing embryos through the newly evolved placenta rather than via the yolk stored in the egg.

In poeciliids, the maternal component of the placenta is derived from the ovarian follicle and called the maternal follicular wall, while the embryonic component of the placenta is the yolk– pericardial sac (henceforth, yolk sac) (15, 16). The independently evolved placentas differ morphologically, ranging from “villous placentas” in which an elaborate maternal follicular wall with villous-like structures interacts with a thin yolk sac, to “smooth placentas”, in which a thin follicular wall meets a more elaborate yolk sac membrane (15, 17, 18). Previous tissue-level gene expression analyses of two independent placenta origins support functional convergence of maternal tissues (19), but much remains unknown, including the identity of the cells that form the independently originated placentas.

The repeated evolution of placentas in poeciliids provides a unique opportunity to address two fundamental questions: (1) What are the molecular, developmental, and evolutionary changes responsible for the origin of a new organ? and (2) How repeatable is evolution — does the convergent evolution of placentas proceed through similar molecular and cellular mechanisms or through diverse ones? We address both questions by studying 9 poeciliid species, encompassing five independent origins of a placenta and their non-placental relatives, using a combination of single-cell transcriptomics and chromatin accessibility, evolutionary genomics, and imaging.

## The cellular basis of independently evolved placentas

We first asked what cell types make up the placenta in *Poeciliopsis turneri*, a species with a placenta characterized by elaborate villous structures on the maternal side (Fig. 1A). To this end, we first assembled and annotated a new chromosome-level reference genome for this species (Supplementary table 1) and then performed single-nucleus RNA sequencing (snRNA-seq) on ovaries from pregnant females (Methods). To better capture ovarian and placental cell populations, we excluded the developing embryos (except for their yolk sacs) from our dissections. The atlas comprised 22,166 cells after quality control (from two libraries; Supplementary figures 1 and 2). We identified all major ovarian cell types based on the expression of known marker genes (Fig. 1B, Supplementary figure 3), including germ cells, the stromal cells that form the matrix supporting the ovary, endothelial cells, immune cells and erythrocytes, germinal epithelial cells, and the cells that make the ovarian follicle – granulosa and theca cells – which support germ cell development and regulate yolk deposition (vitellogenesis). We also identified a single yolk sac cell type.

**Fig. 1.**
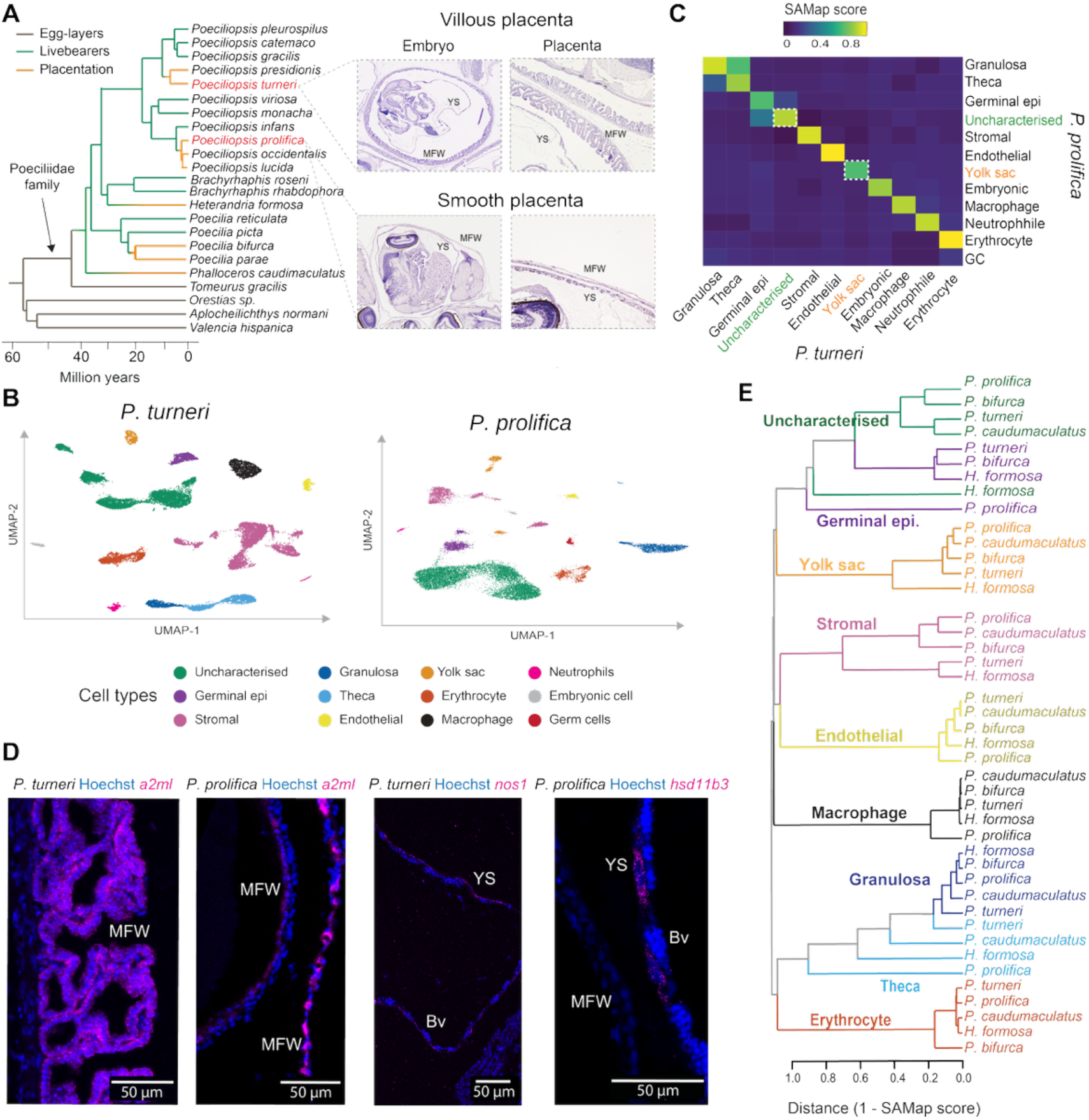
Cellular composition of five independently evolved placentas. **(A)** Phylogenetic tree of the order Cyprinodontiformes and histological sections from *P. turneri* (“villous placenta”) and *P. prolifica* (“smooth placenta”). **(B)** Uniform manifold approximation and projection (UMAP) of the ovarian cells in *P. turneri* and *P. prolifica*. Germinal epi = Germinal epithelium. **(C)** Heatmap showing transcriptome similarity scores (SAMap scores) among ovarian cells of *P. turneri* and *P. prolifica*. **(D)** *In situ* hybridizations on the maternal-embryonic interface of *P. turneri* and *P. prolifica* using markers for the uncharacterized cell (*a2ml*) and the yolk sac cell (*hsd11b3* or *nos1*). MFW = maternal follicular wall, YS = yolk sac, Bv = blood vessels. **(E)** Dendrogram based on distance scores (1 – SAMap score) of all-to-all comparisons of ovarian cells from five placental species.

Unexpectedly, the most abundant cell type in the atlas was the only one whose identity we could not determine based on marker genes or gene ontology analysis of gene expression profiles (Methods). Because this uncharacterized cell population was so abundant, we hypothesized it was contributing to the maternal component of the placenta. To test this hypothesis, we mapped the spatial distribution of this uncharacterized cell type and that of the yolk sac cell (embryonic component of the placenta) within the ovaries of pregnant females using hybridization chain reaction (HCR) with cell-type-specific gene markers (Fig. 1D) (20, 21). These two cell types localized specifically to the maternal–embryonic interface, each forming one of its two tissues: the uncharacterized cell type forms the maternal follicular wall, and the yolk sac cell forms the yolk sac membrane, which is in addition highly vascularized (Fig. 1D). The placenta of *P. turneri* therefore has the simplest cellular architecture possible for a maternal–embryonic interface: one maternal cell type and one embryonic cell type, both associated with an underlying capillary network.

We next asked whether such a simple cellular architecture was shared by independently evolved placentas that additionally had distinct morphologies. We applied the same approach (snRNA-seq of ovarian tissues and chromosome-level genome assembly (Supplementary table 1)) to *Poeciliopsis prolifica*, an independent placental lineage with a smooth-type placenta (Fig. 1A). We annotated the *P. prolifica* ovarian atlas (20,992 cells from three libraries) using the same marker genes as for *P. turneri*. We then compared the transcriptomes of the cell types identified in the two species using SAMap, a cross-species cell-type mapper (22), and distance measures based on cell-type pseudobulks, with both approaches giving similar results (Supplementary figure 4; Methods). These analyses revealed that not only did the placenta of *P. prolifica* have a similarly simple cellular basis as *P. turneri*, but it also comprised the same two cell types, the uncharacterized maternal cell and the yolk sac cell (Fig. 1B,C). This result shows that despite the independent origins of these two placentas and their distinct morphologies, the cells making up this organ are the same in the two species.

To determine whether this shared cellular basis extended to more independently evolved placentas, notably to placentas that evolved in more distant lineages, we generated comparable single-nucleus atlases and genome assemblies for three additional species (Supplementary table 1), *Poecilia bifurca*, *Heterandria formosa* and *Phalloceros caudimaculatus* (20,982, 15,145, and 14,317 cells, respectively), each representing an independent placenta origin (Fig. 1A) (12–14). As before, we annotated cell types within each species using marker genes and compared their transcriptomes across species. These analyses revealed that the ovaries of the five species had the same cell types, including the same maternal and embryonic cells forming the placentas (Fig. 1E). In conclusion, irrespective of placenta morphology and phylogenetic distance, the convergent evolution of placentas in poeciliids has occurred through the repeated deployment of the same two cell types.

## Developmental shift in cell composition of ovarian follicles

Our finding that a yolk sac cell has been repeatedly co-opted to make the embryonic component of the placenta fits within an extensive literature showing yolk sacs have been repeatedly co-opted across vertebrates to make placentas (23, 24). However, our findings for the maternal side were unexpected. Because the maternal follicular wall is derived from the ovarian follicle, we expected it to be made of the granulosa and theca cells that form the follicle. Instead, we found it was made of an undescribed cell type, transcriptomically distinct from granulosa and theca cells, which are also present in our atlases (Fig. 1B,C,E). If this uncharacterized cell type is present in five placental species, parsimony dictates that it must also be present in the phylogenetically intervening non-placental species, where it likely retains an ancestral role. To test this, we generated genome assemblies and performed snRNA-seq on the ovaries from pregnant females of four non-placental species selected to be closely related to each of four placental species (creating four placental–non-placental species pairs; *P. caudimaculatus* was not included due to the absence of a closely related non-placental species) (Fig. 2A). We then performed an all-to-all comparison of cell types using SAMap across all eight species (Fig. 2B). This analysis confirmed our hypothesis: all ovarian cell types are conserved across poeciliids, including the uncharacterized maternal cell type.

**Fig. 2.**
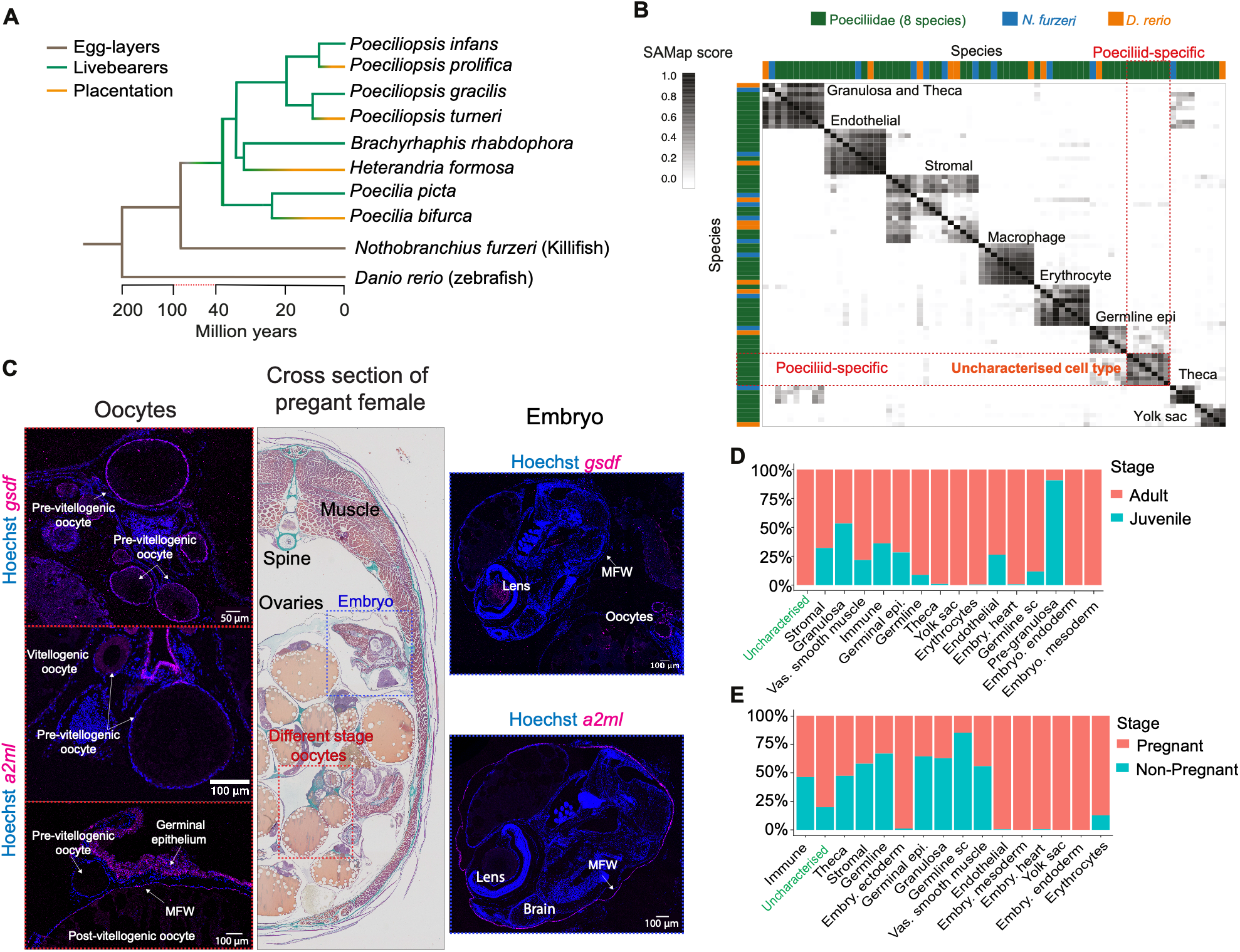
Shift in cell composition of the ovarian follicle. **(A)** Phylogenetic tree with the four pairs of placental–non-placental poeciliids studied and two related egg-laying fish. **(B)** Heatmap of SAMap scores for all ovarian cells (and yolk sac cells in zebrafish) across all species in **(A)**. The red dotted lines highlight the uncharacterized cell type in poeciliids (green bars) and its absence in killifish (blue bars) and zebrafish (orange bars). **(C)** *In situ* hybridization for *gsdf*, a marker of granulosa cells, and *a2ml*, a marker for the uncharacterized cell type, in embryos and oocytes at different stages. **(D)** Cell type proportions in juvenile and adult ovaries of *P. turneri*. **(E)** Cell type proportions in adult pregnant and non-pregnant females of *P. gracilis*.

Next, we investigated the localization of this uncharacterized cell type in non-placental species. Using HCR and the same probe used to spatially locate the maternal cell type in placental species, we found that in non-placental species, this cell type contributes to the ovarian follicle surrounding embryos, similarly to what we had found for placental species (Fig. 2C). Given that granulosa and theca cells are present in all our atlases, this positioning suggested a change in the cellular composition of the ovarian follicle at an earlier developmental stage across all poeciliids. To test for this shift, we localized both granulosa cells and the uncharacterized maternal cells in the ovary of non-placental species using HCR. Because the species in our study can simultaneously carry embryos at different stages of gestation (superfetation), spatially mapping cells also maps them temporally with respect to stages of oogenesis and embryonic development. We found that the follicles surrounding early-stage oocytes up to oocytes accumulating yolk (vitellogenic) are formed by granulosa cells (a conserved feature across vertebrates), whereas late-stage follicles surrounding developing embryos are made of the uncharacterized cell type (Fig. 2C).

To identify when the uncharacterized cell type first emerges during development, we performed snRNA-seq on the ovaries of juveniles of the placental *P. turneri* (6,087 cells) and on the ovaries of non-pregnant adults of the non-placental *Poeciliopsis gracilis* (10,920 cells). Consistent with our imaging, the uncharacterized maternal cell type was absent from juvenile ovaries (Fig. 2D). However, it was already present in the ovaries of non-pregnant adults, surrounding late-stage vitellogenic oocytes (Fig. 2C,E). These results mean that the uncharacterized cells differentiate sometime after vitellogenesis and before fertilization. This observation implies that the differentiation of this cell type is fully under maternal control. Given that the uncharacterized cell appears only in late-stage follicles, henceforth we name it the late follicle cell (LFC).

## Late follicle cells enabled the transition from egg-laying to live-bearing

To understand the functions of late follicle cells, we performed a gene ontology enrichment analysis of their marker genes across all non-placental species. We found conserved enrichments of functional terms suggesting this cell type is epithelial, with strong apical-basal polarity and adherence to its neighbors (Fig. 3A). We then used a set of experimentally validated zebrafish genes (Methods) to investigate the expression of genes associated with two key follicle functions: the regulation of nutrient deposition (vitellogenesis) and egg release (ovulation) in granulosa, theca and late follicle cells (Fig. 3B). Consistent with their roles in other vertebrates, granulosa and theca cells had high expression of all these genes. In contrast, late follicle cells showed expression levels comparable to other non-follicular somatic cell types (Fig. 3B). Together, these analyses suggest that in non-placental species, late follicle cells form an epithelial barrier around late-stage oocytes and developing embryos, and that unlike granulosa cells, they do not contribute to yolk deposition nor respond to ovulation signals. These observations lead to the hypothesis that the shift from granulosa cells to late follicle cells could explain the lack of ovulation in poeciliids and their intrafollicular embryonic development. This cell type could, therefore, have enabled the transition of poeciliids from egg-laying to live-bearing.

**Fig. 3.**
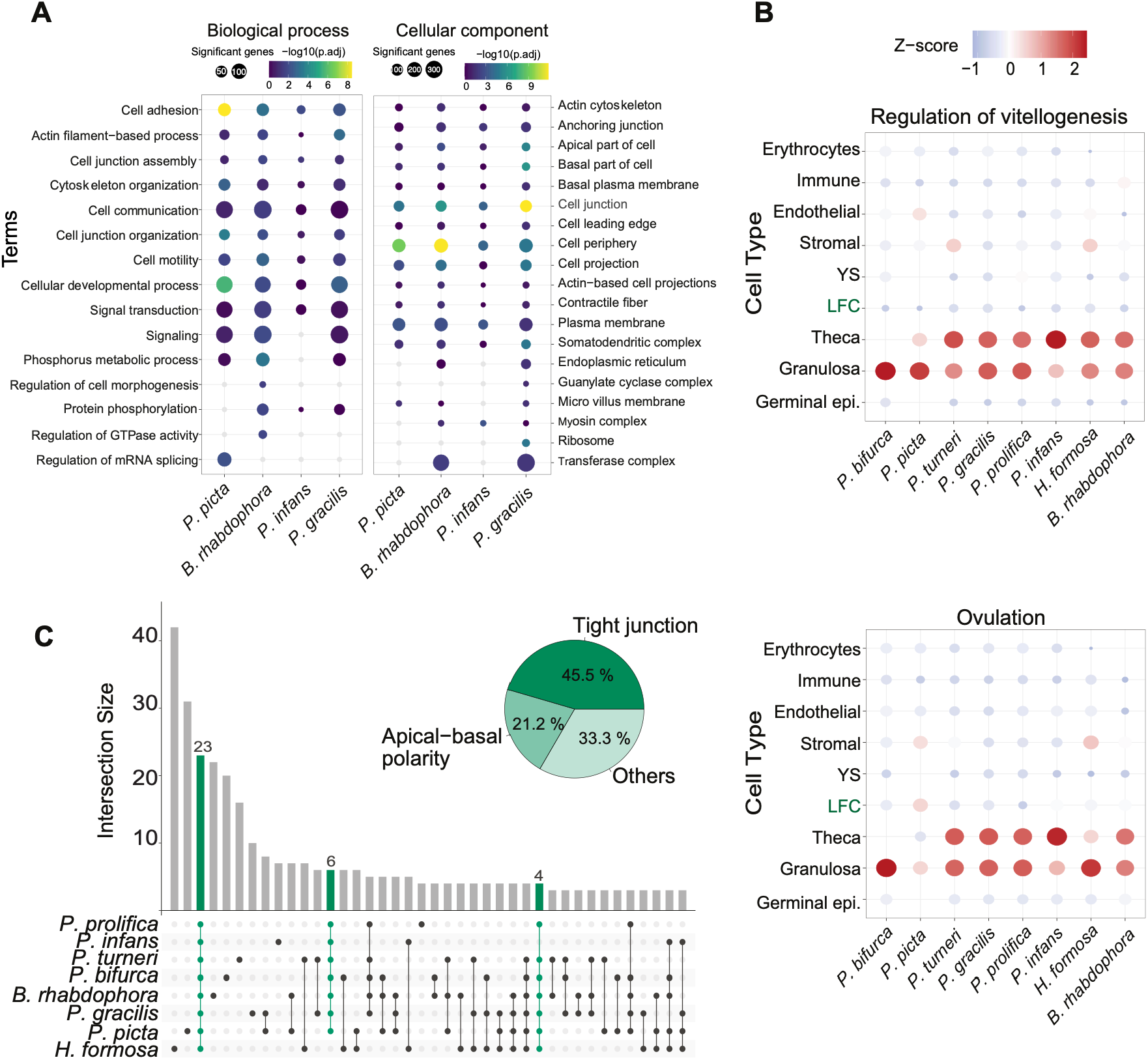
Late follicle cells enabled the evolution of egg-retention. **(A)** Gene ontology analysis of marker genes for late follicle cells of non-placental poeciliids. **(B)** Cell type enrichment (Z-score) for experimentally validated gene sets associated with regulation of vitellogenesis and ovulation across ovarian cell types. **(C)** Overlap across species of genes under positive selection that are highly expressed in late follicle cells. Pie chart shows the gene ontology categories for genes shared across at least seven of the eight species.

If this hypothesis is true, late follicle cells should have evolved in the common ancestor of poeciliids, ∼55 mya (Fig. 2A) (25, 26). To date when this cell type first appeared, we compared our ovarian atlases with publicly available data from other teleosts. We re-analyzed published single-cell RNA-seq datasets of adult zebrafish (*Danio rerio*) and killifish (*Nothobranchius furzeri*) ovaries, as well as Daniocell, a dataset of developing zebrafish embryos (27–30). While zebrafish is a more distantly related teleost, killifish and poeciliids belong to the same order (Cyprinodontiformes) and diverged only ∼75 mya. Using SAMap, we found a one-to-one match for all ovarian cell types across species, except for late follicle cells, which were absent in zebrafish and killifish (Fig. 2B). This result shows that late follicle cells evolved sometime ∼50–75 mya, likely in the ancestor of poeciliids.

Finally, we reasoned that if this new cell type emerged in the ancestor of poeciliids, we might find evidence for positive selection on genes expressed specifically in this cell type across the family, and these genes could further help us understand its evolution. Therefore, we carried out tests for positive selection by inferring rates of protein evolution across the family (for 14,204 1:1 orthologs using PAML, Methods). We detected 1,760 (12%) genes under positive selection, which were enriched for several biological processes, including cell activation, immunity, and response to stimuli, consistent with similar scans in other vertebrate lineages (31–33) (Supplementary figure 5). We next focused on the subset of positively selected genes that are expressed in late follicle cells across multiple species. Because many of these genes are expressed across multiple cell types, we focused on those that are most highly expressed in late follicle cells (Methods, Fig. 3C). This analysis identified 33 positively selected genes highly expressed in late follicle cells in at least seven of the eight species. Strikingly, most of these genes were associated with either tight junctions (15 genes) or apical–basal polarity (7 genes), with the remaining 11 distributed across several other functional categories (Fig. 3C). These results further support the role of late follicle cells in establishing an epithelial barrier at the maternal–embryonic interface and, hence, their role in enabling the transition from egg-laying to egg-retention.

## Late follicle cells evolved through the co-option of an ancestral gene regulatory network

Next, we asked which cell type is the likely progenitor of late follicle cells. Because these cells form a continuous epithelial layer adjacent to late-stage oocytes and the developing embryo, there are two possibilities (Fig. 4A). The first is that late follicle cells differentiate directly from the granulosa cells that form the follicle earlier in development. The second is that late follicle cells differentiate from another cell type that invades/migrates into the follicle before differentiating into late follicle cells (Fig. 4A). While the lack of lineage-tracing tools in poeciliids prevents us from distinguishing between these two hypotheses directly, they make different predictions regarding follicular cell dynamics. If late follicle cells derive from a progenitor population that migrates into the follicle and replaces existing granulosa cells, we should see high levels of cell proliferation (in the progenitor population) and cell death (in granulosa cells). Therefore, we investigated cell proliferation (using PH3) and apoptosis (using Caspase3) in follicle cells before the differentiation of late follicle cells and found few proliferating cells and almost no apoptotic cells (figs. S6A and S6B). An additional challenge to the migration and replacement hypothesis is the presence of a basement membrane between the granulosa and theca cell layers, which constrains cell movement and, when disrupted, leads to ovulation across vertebrates (34, 35). We investigated the presence of the basement membrane during follicle development (using fibronectin) and found it surrounding follicles of all stages, from the early clusters of pre-follicular and oogonia cells (cell nests) through to fertilized follicles (Supplementary figure 6C). Collectively, the low levels of cell proliferation and apoptosis observed in vitellogenic follicles and the persistence of the basement membrane throughout follicle development favor late follicle cells being derived directly from granulosa cells, a type of terminal differentiation. The terminal differentiation of granulosa cells into a phenotypically distinct cell type would not be unprecedented in vertebrates. In mammals, the granulosa cells lying against the basement membrane (i.e., mural granulosa) differentiate after ovulation into the granulosa-lutein cells of the corpus luteum, which are functionally and morphologically distinct (36, 37). If confirmed, this would be an example of convergence between mammals and poeciliids, with both lineages retaining and functionally repurposing granulosa cells to sustain pregnancy.

**Fig. 4.**
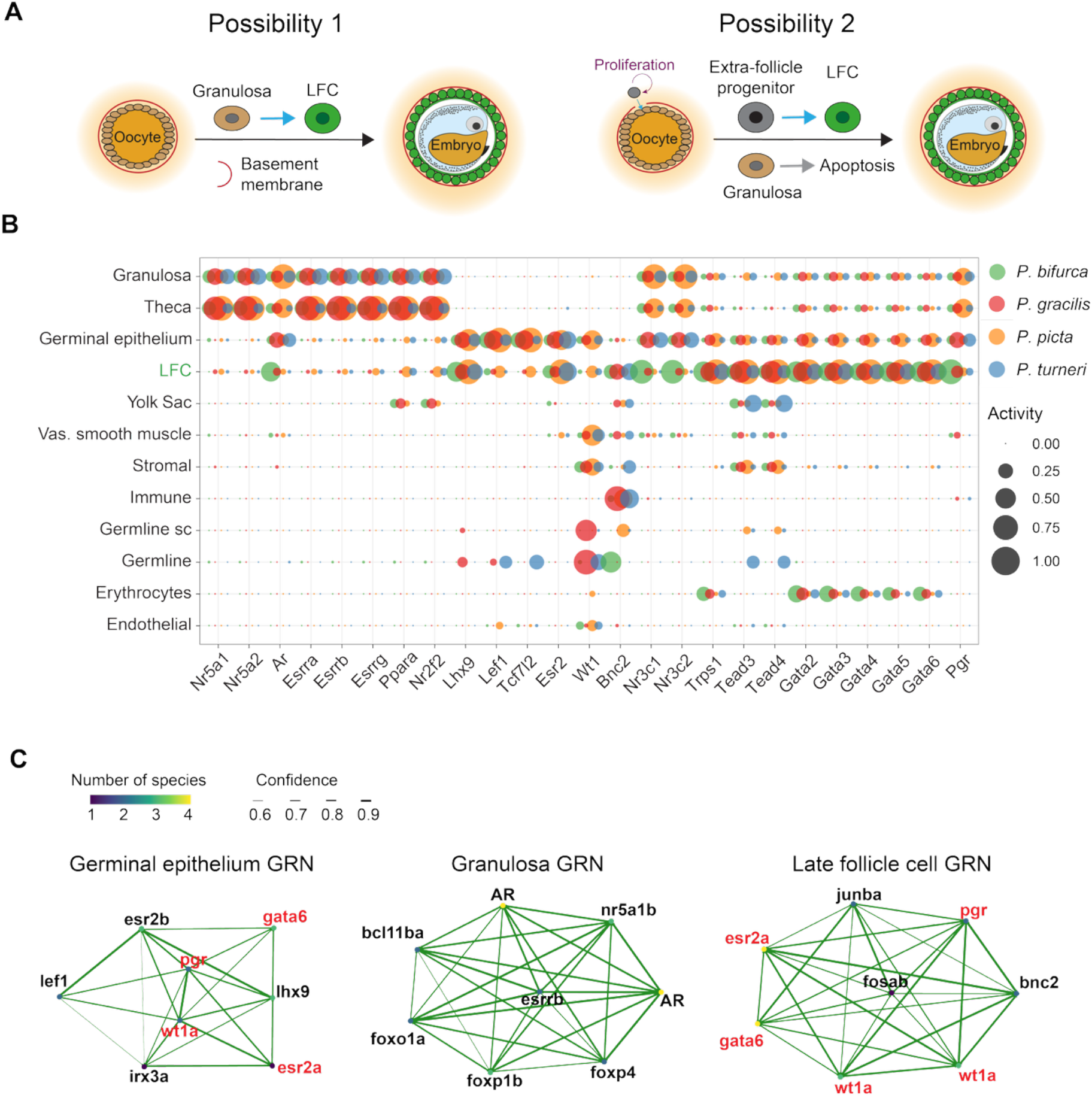
The evolution of late follicle cells. **(A)** Possible developmental origins of late follicle cells. **(B)** TF enrichment in cell type marker peaks of granulosa, theca, germinal epithelium and late follicle cells. **(C)** SCENIC+ reconstructions of the GRNs specifying the germinal epithelium, granulosa cells and late follicle cells. Top 8 regulators per cell type shown. In red are shared regulons between the germinal epithelium and late follicle cell GRNs.

Cell type identity is defined by a gene regulatory network (GRN), and the evolution of a new cell type requires either changes to an ancestral GRN or the emergence of a new one. Therefore, we asked which GRN specifies late follicle cells and how it might have evolved. To this end, we generated chromatin accessibility data by performing single-nucleus ATAC-seq in four species, two placental–non-placental pairs (12,390 cells for *P. turneri*, 12,564 for *P. gracilis*, 5,329 for *P. bifurca* and 8,106 for *Poecilia picta*) (Fig. 4). We first assessed the transcription factor (TF) motifs enriched in the open chromatin of all ovarian cell types (Supplementary figure 7A). This analysis showed that late follicle cells were enriched for estrogen receptor-binding motifs (*esr1*, *esr2*) and gata family motifs (*gata6*), which were also enriched in germinal epithelium cells, the progenitors of granulosa cells (Fig. 4B). This result is consistent with our gene expression analyses, which found germinal epithelium cells to be the most similar transcriptomically to late follicle cells (Fig. 1E, Fig. 2B). The cell-type-defining TF binding sites for late follicle and germinal epithelium cells also show low activity in granulosa cells (Fig. 4B, Supplementary figure 7B). Building on these results, we combined the snRNA-seq and snATAC-seq data for the four species and reconstructed the GRNs of all ovarian cells using SCENIC+ (38). We found the GRNs for the germinal epithelium and granulosa cells are conserved across poeciliids, and recapitulate the known key regulators of these cells in other vertebrates (Fig. 4C; Supplementary figure 7B). The regulons (TFs and downstream genes) in the GRN of the late follicle cell are also conserved across poeciliids (hence, similar between placental and non-placental species) (Fig. 4C). Consistent with the analysis based on chromatin accessibility alone, late follicle cells share five of their eight GRN-defining TFs with germinal epithelium cells, including progesterone receptor (*pgr*), estrogen receptor 2a (*esr2a*) and *wt1a* (Fig. 4C).

Together, our results favor a model in which late follicle cells arise by terminal differentiation of granulosa cells, and in which the new cell type evolved through the partial redeployment (co-option) of a GRN previously active in a progenitor population. The apparent tension between our two lines of evidence — follicular cell dynamics pointing to granulosa cells, and the GRN pointing to germinal epithelium as the immediate cell progenitor — is resolved by noting that in this system, GRN similarity is a poor proxy for developmental relatedness: granulosa cells share none of the core TFs of their progenitor population, the germinal epithelium (39). The resemblance between late follicle cells and germinal epithelium therefore need not reflect direct descent and is better explained by the redeployment of an ancestral GRN in a new cellular context. Definitive resolution will require lineage tracing, which is not yet possible in this family.

## Molecular convergence drove repeated evolution of placentation

Our results show that maternal late follicle cells and embryonic yolk sac cells were independently repurposed five times to form a placenta. We then asked what changed within these cells to enable placentation. The evolution of placentation required modifications to the ancestral roles of both cells: late follicle cells had to change from an epithelial barrier to a surface for nutrient transfer, and the yolk sac cells had to take up nutrients supplied by the mother rather than those stored in the egg. We therefore set out to identify the molecular changes accompanying this shift in each placental lineage and to determine how similar those changes were across independent placenta origins.

We first studied molecular changes in late follicle cells. For four pairs of placental–non-placental species, we identified the genes differentially expressed between the placental species and its non-placental pair and then assigned the direction of expression change based on another non-placental species as an outgroup (Fig. 5A). As expected, considerably more genes changed in gene expression in late follicle cells in placental species than in non-placentals (Fig. 5A), with one exception, the *Brachyrhaphis rhabdophora* – *H. formosa* comparison. An analysis of the differentially expressed genes in the non-placental *B. rhabdophora* showed a significant enrichment for genes involved in gas transport (Supplementary figure 8), which we attribute to a species-specific adaptation of the follicle to larger offspring size (13). We then focused on the genes differentially expressed specifically in the placental species and their biological functions. In three of the four placental species, we found enrichment for the same biological process, lipid metabolism, suggesting functional convergence at the pathway level (Fig. 5B). In contrast, in the placental *H. formosa*, the enrichment was for genes involved in translation and peptide biosynthesis. Notably, all placental species showed an enrichment for genes associated with vesicle- and membrane-related functions, another example of convergence at the pathway level (Fig. 5B). These results are consistent with work based on tissue-level bulk RNA-seq that found an enrichment of lipid metabolism and vesicle-transport in the maternal follicular wall of the placentals *P. prolifica* and *P. presidionis* (a sister placental species of *P. turneri*) (19). Our results also suggest that vesicles are a recurrent feature of placenta evolution, as they have also been proposed to play a role in mammals and placental rockfish (40–43).

**Fig. 5.**
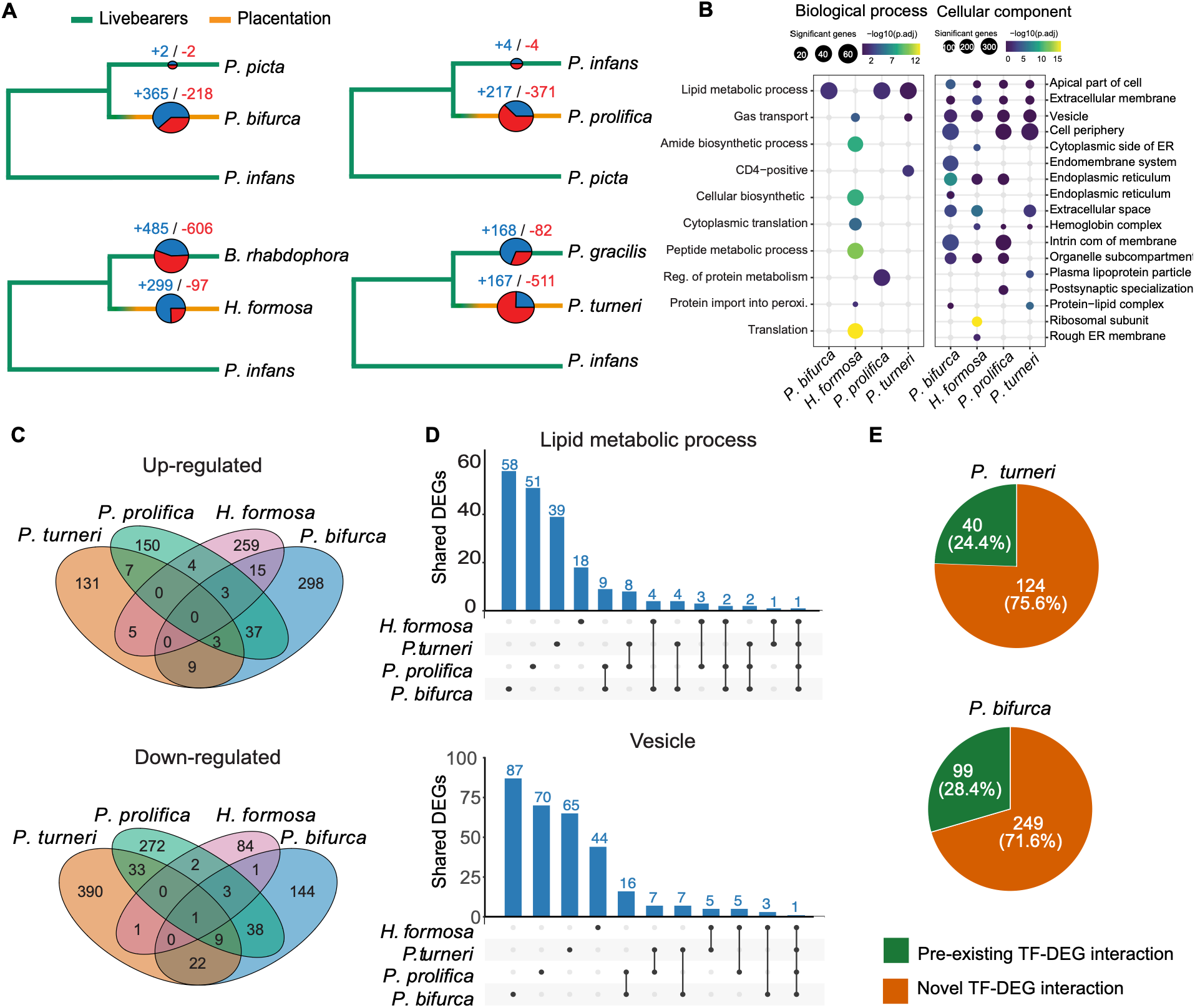
Gene expression changes associated with the evolution of placentation. **(A)** Differentially expressed genes in late follicle cells (blue = up-regulation, red = down-regulation). **(B)** Gene ontology analysis of differentially expressed genes in late follicle cells of placental species. **(C)** Overlap of differentially expressed genes across placental species. **(D)** Proportion of gene sharing across placental species for genes belonging to the gene ontology categories “Lipid metabolic process” and “Vesicle”. **(E)** Proportion of TF-gene interactions involving up-regulated differentially expressed genes (DEGs) in late follicle cells of *P. turneri* and *P. bifurca* that are new or shared with their non-placental pairs.

Given the pathway-level convergence observed in late follicle cells, we asked if this convergence extended to individual genes. We found that most genes that changed expression in placental species were not shared across species (Fig. 5C), a result consistent with the bulk RNA-seq study of the maternal follicular wall of two placental species (19). Although gene sharing was low across species, it occurred more frequently than expected by chance, with some genes being repeatedly recruited during placenta evolution (Supplementary figure 9). *P. bifurca* had the most gene sharing with other placental lineages, while *H. formosa* had the least, consistent with this species being the only placental without an enrichment of genes involved in lipid metabolism (Supplementary figure 9). Furthermore, we examined in detail the genes associated with lipid metabolism and vesicle-related processes, and even within these shared functional pathways, the underlying genes were mostly species-specific (Fig. 5D).

To understand how these differences in gene expression evolved, we compared the GRN reconstructions of the late follicle cells in the four species (two placental–non-placental pairs; Fig. 4C). While the cell-type defining TFs are the same across species, most TF-gene interactions (76% in *P. turneri* and 71% in *P. bifurca*) were unique to the placental species when compared to their non-placental pair (*P. gracilis* and *P. picta*, respectively) (Fig. 5E). This result suggests that the transcriptional changes associated with placentation involved substantial rewiring in the GRN of the late follicle cell type, while keeping the same cell-type defining TFs.

We next studied molecular changes in yolk sac cells. We were surprised to find few total genes changing in expression in the yolk sac (ranging from 58 in *P. turneri* to 85 in *H. formosa*), especially given that considerable numbers of genes changed in expression across all other ovarian cell types (Supplementary figure 10). Because differential gene expression analyses are restricted to genes that are 1:1 orthologs across species, we reasoned we might be missing contributions to yolk sac evolution coming from new genes that duplicated only recently in evolutionary time. Therefore, we studied these genes by estimating when genes first emerged in evolutionary time in each species (using GageTracker (44), Methods). Strikingly, we found an enrichment of new genes specific to poeciliids in the yolk sac of all placental species (Fig. 6A, Supplementary figure 11). This result shows that placentation evolved through different types of molecular changes in the maternal and embryonic components of the placenta. While maternal late follicle cells evolved mostly through changes in gene expression, embryonic yolk sac cells evolved by deploying recently duplicated genes.

**Fig. 6.**
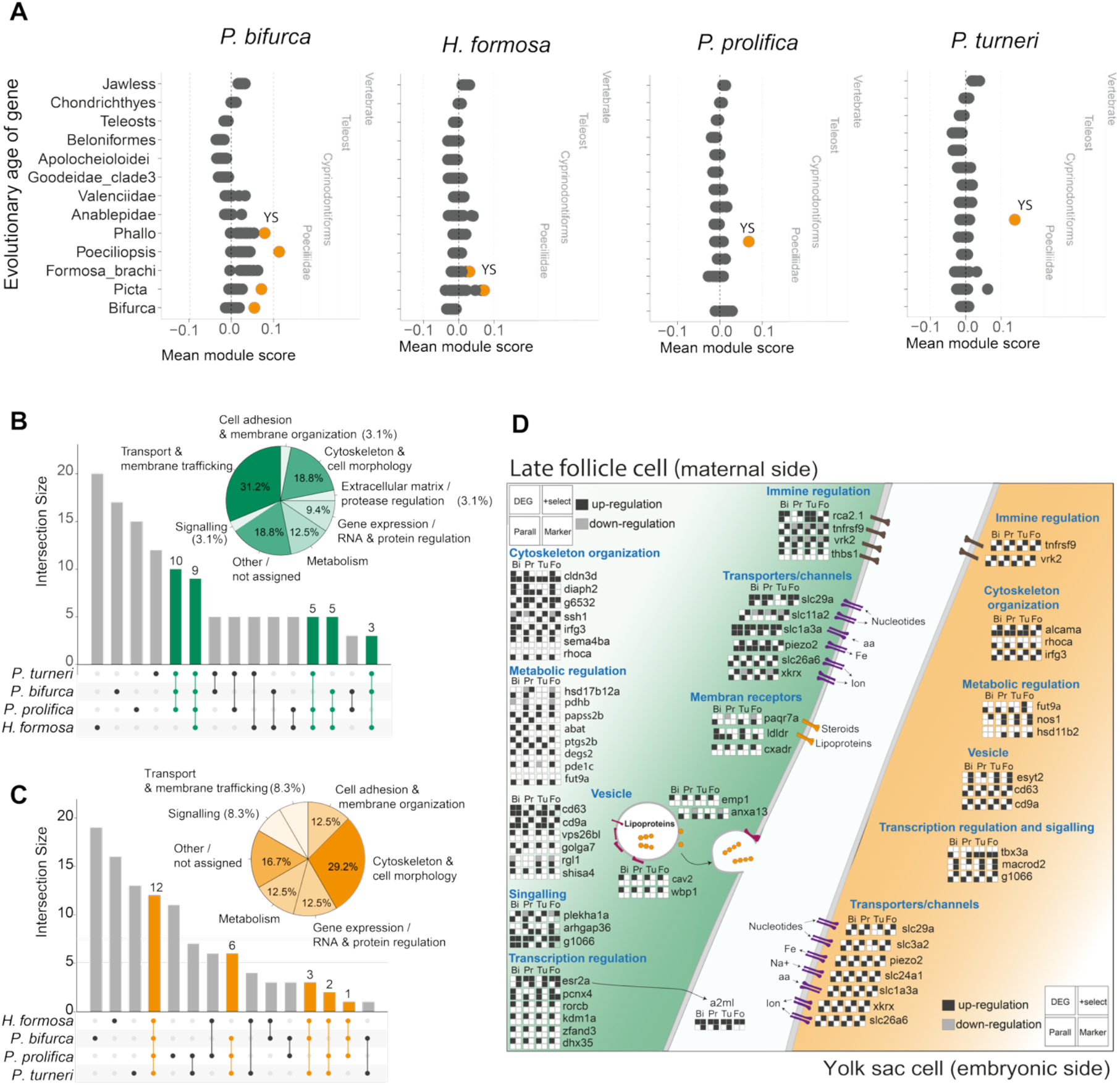
Molecular changes associated with the evolution of placentation. **(A)** Evolutionary gene age distributions across all cell types in four placental species and their relative enrichment to a matched background control. Yellow circles mark the yolk sac cell (YS). **(B** and **C)** Cross-species overlap of genes with parallel substitution expressed in late follicle cells **(B)** or in yolk sac cells **(C)**. The pie charts show the gene ontology categories of genes shared across at least three species. **(D)** Genes with multiple types of molecular change associated with the evolution of placentation in *P. bifurca* (Bi), *P. prolifica* (Pr), *P. turneri* (Tu) and *H. formosa* (Fo). Different types of molecular change are represented by filled (positive) or empty (negative) squares. (DEG = differentially expressed gene in placental species, + select = genes under positive selection, Parall. = genes with parallel substitutions, and Marker = marker genes specific for the cell type).

Given the pathway-level convergence observed at the level of gene expression and the greater than expected gene sharing between placental species, we asked if this convergence also extended to amino acid substitutions. To answer this question, we identified parallel amino acid substitutions that arose independently in placental species (using CSUBST (45), Methods). We defined parallel substitutions as independent substitutions at the same orthologous protein site that resulted in the same amino acid substitution in two or more placental species. We detected a 3.5-fold excess of parallel substitutions in placental species relative to non-placental species: 663 genes with placental-specific parallel substitutions vs 190 genes in non-placental species. Among the 663 genes, 608 have parallel substitutions shared between two placental species (182 in non-placental species), 53 among three species (8 in non-placental species), and two shared across all four placental species (none in non-placental species). We then ranked these genes by expression levels in late follicle cells and yolk sac cells and identified the most highly expressed. In late follicle cells, the number of expressed genes with parallel substitutions ranged from 57 in *P. turneri* to 95 in *P. bifurca*. An analysis of the top 57 (smallest set) most highly expressed genes identified 32 expressed in at least three of the four placental species (Fig. 6B). Most of these genes were associated with transport and membrane trafficking (10 genes; 31%) or cytoskeleton (6 genes; 19%) (Fig. 6B). A similar analysis in the yolk sac identified 24 genes highly expressed in at least three placental species, 7 of which (29%) were associated with the cytoskeleton (Fig. 6C). These results suggest that recurrent parallel substitutions in proteins in placental species contributed to the evolution of nutrient transfer and the re-organization of cell morphology that enabled placentation.

Finally, we combined our analyses of convergence (gene expression and parallel substitutions), positive selection, and cell-type marker genes to identify genes that appeared in at least two analyses in two or more placental species, or in one analysis across all placental species. We found that the same types of genes show changes across multiple placental species (Fig. 6D).

In both late follicle cells and yolk sac cells, these genes are associated with vesicle trafficking, transport and channel activity, metabolism, and cytoskeletal organization (Fig. 6D).

These analyses show that the repeated evolution of placentas involved convergence at every level examined, though to varying degrees. Convergence was strongest at the cell-type level, where the same maternal and embryonic cells were used across all five origins. Functional convergence was also substantial, with lipid metabolism and vesicle-related processes recurring across several placental lineages, alongside parallel changes in genes encoding membrane transporters. By contrast, convergence at the level of individual gene expression was weaker. Although gene sharing between placental species exceeded chance expectations, most expression changes were species-specific, even within shared pathways. Thus, the same epithelial barrier was repeatedly transformed into a nutrient exchange surface through largely different genes. The two halves of the placenta also followed distinct molecular routes, with gene expression changes predominating in maternal late follicle cells and gene duplications in embryonic yolk sac cells.

## Discussion

By studying five independent and recent origins of a placenta in poeciliid fishes, we traced the cellular and molecular paths from egg-laying to live-bearing to placentation. This path unfolded in two steps: first the evolution of a new maternal cell type, which likely enabled the evolution of egg-retention, and then its repeated co-option, together with the embryonic yolk sac, into a placenta.

The transition from egg-laying to egg-retention likely arose in the ancestor of poeciliids by a shift in the cellular composition of the ovarian follicle from granulosa cells to late follicle cells. The molecular profile of late follicle cells—an epithelium enriched for tight-junction and polarity genes, many under positive selection, and lacking the vitellogenesis and ovulation machinery of granulosa cells—can explain both defining features of egg retention: loss of ovulation and embryonic development within the follicle. Because live-bearing is associated with, and may have fueled, a marked increase in poeciliid diversification (8), the origin of this cell type may have been a key innovation (46) that drove the diversification of an entire family.

Late follicle cells are also a rare example of a recently evolved cell type, offering a unique opportunity to investigate how new cell types originate (2). Although lineage tracing is not yet possible in poeciliids, our results are consistent with late follicle cells differentiating from granulosa cells. If true, they are an example of cell-type evolution by terminal addition, in which a terminally differentiated cell type evolves the capacity to differentiate into another cell type (2, 47). Terminal addition has so far been demonstrated empirically only for the origin of melanoleucophores, a white pigment cell in zebrafish (43, 47). Most new cell types have been proposed to originate through the sister cell type model, in which an ancestral multifunctional cell type splits into two descendant cell types that each specialize in a subset of its functions (e.g., ciliary and rhabdomeric photoreceptors, vertebrate rods and cones) (48–51), or through cell type hybridization (or fusion), in which the gene expression programs of two pre-existing cell types are combined in a new single cell (e.g., solvent cells of the rove beetle defensive gland) (50, 52). Late follicle cells suggest terminal addition may be a more common mechanism of cell type origination than currently appreciated.

The GRN of late follicle cells appears to have arisen through partial redeployment of the GRN active in the germinal epithelium, the progenitor of granulosa cells, providing an empirical example of GRN co-option in the origin of a cell type (50, 53, 54). Granulosa cells may be unusually prone to further differentiation. In mammals, after ovulation the ruptured follicle develops into the corpus luteum and granulosa cells differentiate into progesterone-secreting granulosa-lutein cells (55). Furthermore, granulosa cells can transdifferentiate into their male counterpart, Sertoli cells; this can be induced experimentally (56, 57), and in sex-changing fish this granulosa-to-Sertoli cell transition occurs naturally (58, 59). Whether this reflects a general propensity of granulosa cells for further differentiation, or strong selection on the follicle whenever live-bearing evolves, is unknown, but it suggests that repurposing follicle cells may be a recurrent path to viviparity.

We found striking levels of convergence in the transition from egg-retention to placentation. Egg-retention placed a maternal epithelium in direct contact with the embryonic yolk sac, so the evolution of placentation “only” required functional changes in cells that were already in place. This cellular architecture may explain why placentas have evolved repeatedly in poeciliids. Despite their independent origins, these placentas repeatedly modified the same cell types into a functionally similar nutrient exchange surface. This predictability suggests developmental constraints: only a limited set of cellular functions and pathways may be able to mediate nutrient transport and support embryonic development. Yet this shared outcome was achieved through largely lineage-specific molecular changes, suggesting that developmental and functional constraints can channel independent lineages toward similar phenotypes without requiring the repeated use of the same genes. This decrease of convergence from cells to pathways to genes mirrors findings in other systems of parallel adaptation (60, 61). The two halves of the placenta also evolved by different kinds of molecular changes. Maternal late follicle cells evolved largely through changes in gene expression and GRN rewiring, whereas the yolk sac cells showed few expression changes but a marked enrichment of recently duplicated, poeciliid-specific genes, possibly because this cell type was already specialized for nutrient uptake from the yolk (62).

The evolution of new cell types is often thought to provide the basis for the evolution of novel organs by introducing new cellular functions into existing structures. Our findings reveal an alternative route to organ evolution. We show that the evolution of the late follicle cell facilitated egg retention, while subsequent modifications of this cell type enabled the repeated evolution of placentation within the family. Thus, our results reveal a more complex scenario in which recently evolved cell types can provide substrates for further evolutionary adaptation, ultimately facilitating the evolution of a new organ (46).

## Materials and Methods

### Animal husbandry and sample collection

We maintained all poeciliid fish stocks at 26°C (pH ∼7.4, GH ∼180–200) in the Aquatics Facility at the Francis Crick Institute or the vivarium at the University of California, Riverside. We euthanized fish by anesthetic overdose (MS-222 or Aquased), followed by decapitation, in accordance with local animal-use regulations and protocols approved by the relevant institutional ethics committees.

We dissected the maternal follicular wall, yolk sac and whole ovaries from non-pregnant and pregnant females. To isolate the maternal follicular wall, we grasped the tissue with forceps and gently separated the tissue layers to create an opening through which we removed the embryo. We subsequently detached the yolk sac by pinching the yolk sac–embryo boundary with one pair of forceps while gently pulling the embryo away with a second pair. We performed all dissections in PBS or reverse-osmosis (RO) water. For snRNA-seq and snATAC-seq, we snap-froze tissues in liquid nitrogen or on dry ice. For HCR and immunofluorescence, we fixed tissues in 4% PFA. We stored samples for up to two years at −80°C (snRNA-seq and snATAC-seq), −20°C (HCR samples stored in methanol) or 4°C (immunofluorescence).

### Genome assembly and annotation

We generated de novo genome assemblies for *P. gracilis*, *P. turneri*, *P. infans*, *P. prolifica*, *H. formosa*, *B. rhabdophora*, *P. bifurca* and *P. caudimaculatus*. For *P. picta*, we used the previously published genome assembly GCA_033032245.1 (UBC_Ppic_1.0) (63). We extracted genomic DNA from muscle tissue using the Blood & Cell Culture DNA kit (QIAGEN, cat. no. 13343). We generated draft genome assemblies from Oxford Nanopore long reads sequenced on a PromethION P2 platform (Oxford Nanopore Technologies, cat. no. PRO-SEQ002) using Flye (64). We performed error correction and consensus generation with Racon (65), followed by additional polishing with short-read DNA sequencing data generated from the same sample using POLCA (66). For all species except *B. rhabdophora*, we scaffolded the draft assemblies to chromosome level using Hi-C data generated from muscle tissue with the Arima High Coverage Hi-C Kit and the Arima mapping pipeline (https://github.com/ArimaGenomics/mapping_pipeline/). We assessed genome assembly quality using BUSCO completeness scores with the Cyprinodontiformes lineage dataset and assembly contiguity using the N50 statistic (67).

We identified repetitive elements in all genomes using RepeatModeler v2.0 and subsequently soft-masked the genomes using RepeatMasker v4.1.3 (68, 69). Following repeat masking, we annotated the genomes using BRAKER3 (70). We incorporated protein evidence from the vertebrate OrthoDB database v12 (71) (as recommended by BRAKER3), and bulk RNA-seq data from brain, ovarian follicles, and whole-body tissues as extrinsic evidence to support gene structure annotation. We annotated ribosomal RNA (rRNA) genes using Barrnap with the ‘--kingdom euk’ parameter (72) and annotated mitochondrial genomes using MitoFinder (73).

To improve snRNA-seq read mapping, we extended the 3′ UTR annotations of coding genes using Funannotate v1.8.15 (74), specifically its ‘updatè function. This function validates gene models, recovers missing genes and annotates UTRs using bulk RNA-seq evidence. We subsequently refined the 3′ UTR annotations using peaks2utr v1.2.0 (75), which identifies strand-specific transcription termination sites from aligned snRNA-seq reads and extends or annotates 3′ UTRs based on candidate peaks assigned to the nearest gene.

### Gene ontology annotation

We assigned Gene Ontology (GO) terms to all annotated genes in each genome using FANTASIA v4 (76). FANTASIA predicts GO annotations by embedding protein sequences using the ProtT5 protein language model and transferring annotations from experimentally characterized reference proteins based on embedding-space similarity (GOPredSim). For each species, we used the predicted proteome as input to FANTASIA, removed identical protein sequences and excluded sequences longer than 5,000 amino acids, following the default FANTASIA pipeline settings. We ran FANTASIA independently for each of the nine species included in this study.

### Gene orthology annotation

We grouped annotated genes into orthogroups using the pipeline described by Bontonou et al. (77). For each species, we translated coding sequences into peptide sequences and retained the longest transcript for each gene using AGAT v1.0 (78). We combined the resulting peptide sequences from all species into a single directory and used these as input for OrthoFinder v2.5.4 (79), with the -M msa option to infer gene trees. We further resolved non-1:1 orthology relationships identified by OrthoFinder using Possvm v1.0 (80). We aligned protein sequences using MAFFT v7.526 with the --auto option (81) and inferred maximum-likelihood phylogenies with bootstrap support using IQ-TREE v1.6.12 (82). Possvm then refined the OrthoFinder orthogroups by parsing the resulting gene trees using a species-overlap algorithm and clustering the sequences with the Markov Cluster Algorithm (MCL). This procedure identified 2,382 additional 1:1 orthologous genes relative to the 11,822 initially identified by OrthoFinder.

### Generation of snRNA-seq datasets

We isolated nuclei from flash-frozen tissues by pestle homogenization, adapting published protocols (83, 84). Because fish nuclei are smaller than mammalian nuclei, we increased the centrifugation speed and duration to improve nuclear recovery and incorporated RNase inhibitors with activity against fish RNases. We homogenized tissue using a 1.5-ml Eppendorf pestle (SP Bel-Art, cat. no. 199230001) in ice-cold homogenization buffer (HB) containing 250 mM sucrose, 25 mM KCl, 5 mM MgCl₂, 10 mM Tris-HCl (pH 8.0), 1 µM DTT, 0.4 U µl⁻¹ RNasin Plus (Promega, cat. no. N2611), 0.2 U µl⁻¹ SUPERase-In (Invitrogen, cat. no. AM2696), 0.4 U µl⁻¹ Ambion RNase inhibitor (Invitrogen, cat. no. AM2684) and 1× protease inhibitor cocktail (Roche, cat. no. 11836170001). We cleared the homogenate of debris by centrifugation at 100g for 1 min twice, and washed the resulting nuclear pellet 2–3 times in HB by centrifugation at 600g for 6 min at 4°C. We resuspended nuclei in ice-cold storage buffer containing 166.5 mM sucrose, 5 mM MgCl₂, 10 mM Tris-HCl (pH 8.0), 0.2 U µl⁻¹ SUPERase-In, 0.4 U µl⁻¹ RNasin and 0.4 U µl⁻¹ Ambion RNase inhibitor, and passed the suspension through a 30-µm filter (Miltenyi Biotec, cat. no. 130-098-458).

All snRNA-seq samples except for SAF4933A57 (the juvenile P. turneri sample) were prepared according to this method. For SAF4933A57, the nuclei were isolated according to the method used for snATAC-seq samples instead, with the addition of equivalent concentrations of RNase inhibitors to the buffer because this protocol performed better with low amounts of tissue input.

We generated libraries using the 10x Genomics Chromium Single Cell 3′ v3.1 RNA assay according to the manufacturer’s protocol (85). Before loading, we stained nuclei with propidium iodide/acridine orange (Sciquip, cat. no. F23001) and counted them using a LUNA-FX7 automated cell counter (Logos Biosystems, cat. no. L70001). We loaded 22,000 nuclei per lane, targeting ∼10,000 recovered nuclei, while empirically adjusting the loading concentration to account for the smaller size of fish nuclei. Library preparation was done according to the manufacturer’s protocol. We sequenced libraries on an Illumina NovaSeq 6000.

### snRNA-seq data processing and quality control

We generated genome indices using the genomeGenerate function in STAR and mapped and quantified libraries against these references using STARsolo v2.7.10b (86). We performed alignment using the alignReads command with default parameters, except for outFilterScoreMin, which we set to 20. We resolved multimapping reads using the expectation–maximization algorithm implemented in STARsolo.

We retained nuclei-containing droplets using an approach that combines the estimated fraction of intronic reads, which will be higher in nuclei than in empty droplets containing ambient RNA (“soup”), with the total number of uniquely mapped reads (UMIs). We determined thresholds for both metrics independently for each library by manually inspecting the resulting plots (87). We removed ambient RNA using SoupX v1.5.2 (88). We generated a contamination profile for each library using estimateSoup, quantified the contamination fraction using autoEstCont and corrected the expression counts in nuclei-containing droplets using adjustCounts.

We imported the corrected count matrices into Seurat v5.4.0 for further quality-control filtering (89). We removed droplets in which more than 8% of reads mapped to ribosomal RNAs or ribosomal proteins. We then removed ribosomal genes from the count matrix to avoid biasing downstream clustering. Because sequencing depth varied between libraries, we defined library-specific thresholds for the minimum numbers of UMIs and detected features based on their respective count distributions, while requiring at least 750 UMIs and 500 features in all libraries. We also excluded droplets with unusually high numbers of UMIs (>10,000) or detected features (>3,000), which were likely to represent doublets.

We identified putative doublets using DoubletFinder v2.0.4 (90). We optimized the neighborhood size parameter (pK) for each library according to the developer’s recommended procedure and assumed a 10% expected doublet rate, adjusted for the estimated proportion of homotypic doublets, following 10x Genomics guidelines. We removed cells classified as doublets, as well as clusters in which more than 50% of cells were classified as doublets.

### Clustering and annotation

We processed the high-quality nuclei using the standard Seurat workflow. We normalized counts using LogNormalize with a scale factor of 10,000 (91). We identified the 5,000 most highly variable genes (HVGs) using FindVariableFeatures with the vst method and scaled the data using ScaleData. We performed principal component analysis (PCA) on the HVGs using ‘RunPCÀ. We constructed a k-nearest-neighbor graph using ‘FindNeighbors’ over library-specific principal components 1–30 and identified clusters using Louvain graph-based clustering with FindClusters at a resolution of 0.3 (92). We visualized the final clusters using UMAP (‘RunUMAP’) based on principal components 1–20 (91).

We manually annotated clusters in each library by comparing their marker-gene profiles with published zebrafish single-cell datasets spanning ovarian and embryonic cell types (28–30, 93). We identified cluster marker genes using FindAllMarkers with a Wilcoxon rank-sum test and the following parameters: min.pct = 0.25, logfc.threshold = 0.25 and min.diff.pct = 0.3. We compared marker-gene profiles with those reported in the reference datasets and with additional evidence from the literature, including *in situ* hybridization and immunofluorescence staining (94).

We integrated all libraries from each species to generate species-level atlases using Harmony (pregnant datasets) and scVI (juvenile and non-pregnant datasets) (95, 96). We performed normalization, scaling and feature selection on the raw counts of the integrated objects. We constructed a k-nearest-neighbor graph using the integrated latent space with FindNeighbors (dims = 1:25) and identified clusters using Louvain community detection. We evaluated a range of clustering resolutions for each species and selected the resolution that provided the optimal separation of clusters for cell-type annotation. We visualized the integrated datasets using UMAP (RunUMAP, dims = 1:25). We annotated clusters as described above.

### Cross-species comparisons using pseudobulks

To obtain cell-type-specific transcriptomes, we generated pseudobulk profiles for each species atlas using the AverageExpression function in Seurat. We standardized expression values across species by calculating gene-wise Z-scores across all matched cell types using the scale function in base R. This transformation centered the expression of each gene at zero and scaled it to a standard deviation of one, thereby placing genes on a comparable scale and reducing the influence of differences in absolute expression levels between species. We restricted subsequent analyses to 1:1 orthologs across species analyzed. We calculated pairwise Pearson correlation coefficients between pseudobulk transcriptomes using the correlate function from the corrr R package v0.4.5 (97).

### Cross-species integration with SAMap

To integrate single-nucleus RNA-sequencing profiles across species, we used SAMap (22). We converted the Seurat objects to AnnData objects and generated pairwise gene mappings between species using BLAST v2.14.0 (98), following the developers’ recommended parameters (outfmt 6, max_hsps 1 and an E-value threshold of 1 × 10⁻⁶), adapted from the map_genes.sh script provided with SAMap. We ran SAMap using the sm.run function with default parameters and identified cell–cell similarities using a mapping-score threshold of 0.2 (hom_edge_thr = 0.2). We extracted the resulting cell–cell mapping scores using get_mapping_scores for downstream analyses.

### Cell-type tree construction

We constructed cell-type trees using two complementary measures of cell-type similarity. First, we quantified transcriptomic similarity using pseudobulk gene expression profiles normalized by a gene-specificity index (GSI), following Tosches et al. (99). For each species, we calculated the mean expression of each gene across cell types and divided its expression in each cell type by this mean, thereby expressing gene expression relative to its average across cell types. We then calculated pairwise Pearson correlation coefficients between cell-type profiles across genes and converted these coefficients to distances using (d = 1-r). To assess the robustness of the inferred relationships, we generated 1,000 bootstrap replicates by resampling genes with replacement, recalculating the correlation matrix and reconstructing the tree for each replicate (87). Node support was calculated as the proportion of bootstrap trees containing each clade.

We also constructed cell-type dendrograms based on the cross-species cell-type scoring inferred by SAMap (22). We extracted the cell-type mapping-score matrix using get_mapping_scores and converted mapping scores to distances using 1-SAMap mapping score. We performed hierarchical clustering using hclust with Ward.D2 linkage and visualized the resulting trees as dendrograms.

### Cross-species differential expression analysis

We identified genes differentially expressed between placental and non-placental species within matched cell types using a pairwise cross-species analysis. To minimize species-specific differences in read alignment and gene annotation, we first filtered sequencing reads using the CrossFilt pipeline (100), applying the standard pipeline independently to each species pair. We subsequently remapped the filtered reads to the corresponding species-specific reference genomes and quantified gene-level expression using STARsolo v2.7.10b (86).

For each species pair and homologous cell type, we generated pseudobulk count matrices using the function AggregateExpression from Seurat across nuclei within each biological sample. We restricted the analysis to 1:1 orthologs and retained genes detected in both species. We used the resulting matched orthologous genes as the basis for cross-species comparisons, assigning the gene identifier of one reference species to each orthologous pair to facilitate comparisons across species. We then performed differential expression analysis using DESeq2 on the raw pseudobulk counts (101). Libraries from a given species were treated as independent replicates. We fitted a negative-binomial generalized linear model and estimated differential expression between the two species in each pair. We applied library-size normalization and estimated gene-wise dispersions using the standard DESeq2 workflow. We shrank estimated log2 fold changes using the adaptive shrinkage method (ashr) to improve the stability of effect-size estimates for genes with low counts or high dispersion. We considered genes differentially expressed when they showed an absolute log2 fold change of ≥1.5 and an adjusted P value of ≤0.05.

To identify lineage-associated expression differences, we compared the direction and magnitude of differential expression across three pairwise species comparisons. For each focal placental lineage, we required the focal species to show a consistent expression difference relative to both its non-placental counterpart and an additional non-placental outgroup species. We classified a gene as upregulated in the focal lineage when its log2 fold change was ≥1.5 in both comparisons, and both comparisons had an adjusted P value of ≤0.05. Conversely, we classified a gene as downregulated when its log2 fold change was ≤−1.5 in both comparisons with an adjusted P value of ≤0.05.

To test whether lineage-specific differentially expressed genes (DEGs) were shared between species more frequently than expected by chance, we compared DEG sets across species using 1:1 orthologs. For each species pair, we restricted both DEG sets and their corresponding gene universes to the shared 1:1 ortholog universe and calculated the observed number of overlapping genes. We generated null distributions by randomly sampling, without replacement, the same number of genes as observed in each DEG set from the shared orthologue universe. We repeated this procedure 10,000 times and calculated an empirical P value as the proportion of permutations yielding an overlap equal to or greater than the observed overlap. We applied the same approach to assess the overlap shared across three and four species. We separately assessed whether shared DEGs showed consistent direction of expression by comparing the direction of differential expression for each shared ortholog across species. Genes were considered directionally concordant when all species showed the same direction of change.

### Gene Ontology enrichment analysis

We tested for GO terms using topGO (102). For each gene list analyzed, we defined the gene universe as all genes included in the corresponding table. We restricted the target set to genes with GO annotations available from FANTASIA and used the same GO annotation set to construct gene-to-GO mappings. We performed enrichment analyses separately for the three GO ontologies: biological process (BP), molecular function (MF) and cellular component (CC). We used the topGOdata function with the annFUN.gene2GO annotation function and tested GO-term enrichment using Fisher’s exact test with the parentchild algorithm. We extracted enrichment statistics for all GO terms tested and corrected the resulting P values for multiple testing using the Benjamini–Hochberg procedure. We considered GO terms with an adjusted P value <0.05 to be significantly enriched.

### Hybridization chain reaction (HCR)

We adapted the HCR protocol used in this study from previous literature (20, 21, 103, 104). We fixed ovaries in 4% paraformaldehyde (Thermo Scientific Cat# 28908) overnight at 4°C, dehydrated them through a graded methanol series, and stored them in 100% methanol at −20°C. For sectioned material, we paraffin-embedded the samples, cut them at 6 µm onto slides (Fisher Scientific Cat# 10149870), dried them overnight at 37°C, and stored them at room temperature. Sequences of HCR probes used in this study are available in Supplementary table 2.

### Paraffin sections

We baked slides (65°C, 30 min) and deparaffinized them in xylene (Merck Cat# 534056; 2 × 4 min) followed by 100% ethanol (3 × 4 min). We permeabilized sections with proteinase K (1 µl of 20 mg/ml stock, Thermo Scientific Cat# EO0491, in 3 ml PBS) for 5 min at 37°C and pre-hybridized them in hybridization buffer (Molecular Instruments) for 30 min at 37°C. We applied probe solution (4 pmol probe per 100 µl hybridization buffer; ∼25 µl per section under a hybrislip) overnight at 37°C. We then removed probes through a graded probe-wash-buffer/SSCT series (Molecular Instruments; Thermo Scientific Cat# AM9770) at 100/75/50/25/0% (5 × 15 min) with a final 15-min wash in 5% SSCT at room temperature.

We equilibrated sections in amplification buffer (30 min, RT) while separately snap-cooling the H1 and H2 hairpins; we then applied the hairpins at 60 nM (2 µl of 3 µM stock per 100 µl amplification buffer) overnight in the dark at room temperature. We used the B1-647 hairpin for all images. Next, we removed excess hairpins (3 × 10 min, 5% SSCT), stained nuclei with 1:500 Hoechst (Thermo Scientific Cat# 62249) for 20 min, and washed the sections (3 × 5 min SSCT). Finally, we mounted slides in SlowFade Gold Antifade Mountant (Fisher Scientific Cat# 11559306) under a coverslip (Dixon Glass Cat# NPS13/2460) and imaged them on a Zeiss 710 or Leica SP8 confocal microscope.

### Whole-mount juvenile ovary

We followed the HCR protocol described above with several modifications. We rehydrated samples through a graded methanol/PBST series and incubated them with proteinase K for 45 min at room temperature. We extended the pre-hybridization step to ≥1 h and performed four 15-min washes in 100% probe-wash buffer. Before the final SSCT washes, we stained nuclei with Hoechst (1:1,000) for 2 h. Finally, we mounted samples in 0.1% low-melting-point agarose (Sigma-Aldrich Cat# A4018) in glass-bottom dishes (Thermo Fisher Scientific Nunc Cat# 150682) and acquired images using a Leica SP8 confocal microscope. We provide probe sequences and working concentrations in Supplementary table 2.

### Immunofluorescence

We adapted the immunofluorescence protocol from a method developed for staining zebrafish ovaries (105). We processed, embedded, sectioned and deparaffinized paraffin sections as described above for HCR. We performed antigen retrieval by boiling the sections in citrate buffer (Cell Signaling Technology, cat. no. 14746) for 10 min, followed by cooling to room temperature. After three washes in water, we blocked the sections for 1 h in PBS containing 0.3% Triton X-100, 10% fetal bovine serum (Gibco, cat. no. A5209401), 5% donkey serum (Merck, cat. no. D9663) and 10% BSA (Thermo Scientific, cat. no. 37520). We incubated sections with primary antibodies overnight at 4°C (Supplementary table 3). We washed the sections four times in PBS containing 0.1% Triton X-100 (Thermo Scientific, cat. no. A16046.AE), incubated them with secondary antibodies and Hoechst for 1 h at room temperature, and performed three additional washes. We mounted the sections in SlowFade Gold Antifade Mountant (Fisher Scientific, cat. no. 11559306) under coverslips (Dixon Glass, cat. no. NPS13/2460) and acquired images using Zeiss 710 or Leica SP8 confocal microscopes.

### Gene-set module scoring

To quantify the transcriptional representation of predefined gene sets across species, we calculated module scores using the AddModuleScore function in Seurat. We selected genes based on experimental evidence from zebrafish studies demonstrating their involvement in ovulation (https://zfin.org/GO:0030728; Li and Ge, 2020) or the regulation of vitellogenesis (106). We applied the same gene sets independently to each species atlas using the corresponding species-specific gene annotations. The ovulation gene set comprised pgr, inhbb, inhba, fshr, lhcgr, ptgs2, adamts1, cyp17a2, duox, foxp3a, inha, lepr, lhb, mettl3, nedd8 and pgrmc1. The vitellogenesis-regulation gene set comprised fshr, lhcgr, lhr, bmpr2, alk, npr2, fabp, stra6, rbp4, slc7a5, slc38a3, glut1, slc2a1, cyp19a1a, hsd3b, cyp17a1, gja1 and gja4. We additionally calculated module scores for sets of genes assigned by GageTracker to different evolutionary age categories (see below), using the same procedure. This analysis allowed us to assess the transcriptional representation of genes that emerged at different points during evolutionary history and compare their distribution across cell types and species.

### Generation of snATAC-seq datasets

We placed flash-frozen tissue in a Miltenyi C Tube (cat. no. 130-093-237) containing 2 mL nuclei extraction buffer (Miltenyi, cat. no. 130-132-997) supplemented with 0.1% Nonidet P-40 substitute (Thermo Scientific, cat. no. J19628.K2), 0.2 U µl⁻¹ SUPERase-In, 0.4 U µl⁻¹ RNasin and 0.4 U µl⁻¹ Ambion RNase inhibitor. We dissociated the tissue for 5 min using a gentleMACS OCTO homogenizer (Miltenyi, cat. no. 130-134-029) fitted with an OCTO Cooler (Miltenyi, cat. no. 130-130-533). We transferred the homogenate to a 15-ml tube, rinsed the C Tube with an additional 2 mL of extraction buffer and pooled the wash with the homogenate before passing the suspension through a 70-µm filter (Miltenyi, cat. no. 130-098-462). We pelleted nuclei by centrifugation at 500g for 5 min at 4°C and resuspended them in 1 mL resuspension buffer containing PBS, 14% Miltenyi extraction buffer, 1% BSA, 0.2 U µl⁻¹ SUPERase-In, 0.4 U µl⁻¹ RNasin and 0.4 U µl⁻¹ Ambion RNase inhibitor. We passed the suspension through a 30-µm filter and repeated the centrifugation step (500g, 5 min, 4°C). We then resuspended the nuclei in 450 µl ice-cold resuspension buffer, added 50 µl anti-nucleus microbeads (Miltenyi, cat. no. 130-132-997) and incubated the suspension at 4°C for 15 min.

We applied the bead-labeled nuclei to a magnetic LS column (Miltenyi, cat. no. 130-042-401; stand, cat. nos. 130-091-051 and 130-042-303) and washed the column three times with 1 mL resuspension buffer. We removed the column from the magnet and eluted the retained nuclei by applying 1 mL resuspension buffer with the plunger. Following a final centrifugation step (500g, 5 min, 4°C), we resuspended the nuclei in 200–300 µl ice-cold 1× Nuclei Buffer (10x Genomics, cat. no. PN-2000153).

We prepared libraries using the 10x Genomics Chromium Single Cell ATAC v2 assay according to the manufacturer’s protocol (107). We counted nuclei as described above and diluted them to ∼5,000 nuclei µL⁻¹ before loading 22,000 nuclei into the transposition reaction per lane, targeting ∼10,000 recovered nuclei. We sequenced libraries on an Illumina NovaSeq 6000.

### snATAC-seq data processing and quality control

We processed the snATAC-seq libraries with Cell Ranger ATAC v2.1.0 using the count function with default parameters (108). We generated custom reference genomes for *P. gracilis*, *P. turneri*, *P. picta* and *P. bifurca*. For each species, we restricted the GTF annotation to protein-coding transcripts and defined gene boundaries as the minimum and maximum coordinates across all transcripts associated with each gene. We extracted chromosome-level scaffolds from the genome assembly using SAMtools faidx v1.18 (109) and converted the resulting FASTA files to 2bit format using UCSC faToTwoBit v482 (110). We used forgeBSgenomeDataPkg from BSgenome v1.66.3 (111, 112) with species-specific seed files to generate and install custom BSgenome packages, which we then supplied to ArchR createGenomeAnnotation (112).

We generated species-specific ArchR gene annotations from the filtered GTF files. We constructed TxDb objects using makeTxDbFromGFF from GenomicFeatures v1.50.4 (111) and derived transcription start sites (TSSs) from transcript ranges using promoter coordinates. Gene ranges were generated by reducing transcript ranges by gene_id, and exon ranges were imported directly from the GTF. We combined TSS, exon and gene ranges using ArchR createGeneAnnotation and used the resulting annotations for Arrow-file generation and gene-score calculation. To prevent errors in gene-score calculation at contig boundaries, we removed genes whose start coordinate was within 7 kb of a chromosome end, together with their associated TSS and exon annotations.

We configured ArchR v1.0.3 for non-model genomes by disabling chromosome-prefix enforcement (addArchRChrPrefix = FALSE) (112). We imported fragment files using createArrowFiles, retaining cells with TSS enrichment ≥4 and 1,000–20,000 unique fragments (minTSS = 4, minFrags = 1,000, maxFrags = 20,000). We generated a 500-bp genome-wide tile matrix and gene-score matrix during Arrow-file creation. We assessed data quality from per-cell TSS-enrichment and log10(fragment-count) distributions, fragment-length periodicity, TSS-enrichment profiles, peak-type composition of union peak sets and the number of reproducible peaks detected per cell type.

We identified putative doublets using ArchR addDoubletScores with k = 10, knnMethod = “UMAP” and nTrials = 5, using LSI on 1,000 variable features and 20 dimensions. We removed predicted doublets using filterDoublets with cutEnrich = 1.5, cutScore = -Inf and filterRatio = 3.

### Clustering and annotation

We performed iterative latent semantic indexing (LSI) on the tile matrix using ArchR addIterativeLSI with four iterations, 25,000 variable features, 30 dimensions, a clustering resolution of 0.2 and 10,000 cells sampled per iteration. We generated UMAP embeddings using 15 nearest neighbors, a minimum distance of 0.6 and cosine distance (113), and t-SNE embeddings using a perplexity of 30 (114). We identified clusters using the Seurat graph-based clustering method implemented in ArchR (addClusters; resolution = 0.5). For *P. gracilis* and *P. turneri*, which were represented by multiple libraries, we combined samples and corrected batch effects using Harmony (addHarmony) with sample identity as the grouping variable. *P. picta* and *P. bifurca* were each represented by a single library and therefore did not require batch correction.

We identified cluster marker genes from the gene-score matrix using ArchR getMarkerFeatures with a Wilcoxon test, correcting for TSS enrichment and log10(fragment number), and retaining markers with FDR ≤0.01 and log2 fold change (log2FC) ≥1. We visualized gene activity on UMAP embeddings using MAGIC imputation (addImputeWeights) (115). We used accessibility of established marker genes to assign initial cluster identities and subsequently integrated the snATAC-seq data with the corresponding species-specific snRNA-seq atlas to establish final cell-type annotations.

We first performed an unconstrained integration after reclustering the snATAC-seq data at high resolution. We then performed a constrained integration to resolve discrepancies identified during the unconstrained integration by restricting ATAC clusters and RNA cells to matched cell-type groups. We retained the constrained gene-integration matrix in the Arrow files (addToArrow = TRUE) and assigned each ATAC cluster the cell-type label corresponding to the most frequent predicted group.

### Peak calling and annotation

We generated pseudobulk replicates for each cell type using ArchR addGroupCoverages. We identified reproducible peaks using MACS2 v2.2.9.1 through addReproduciblePeakSet, grouping cells by cell type and specifying an effective genome size equal to the number of non-ambiguous bases in the corresponding genome assembly (112, 116, 117). We combined reproducible peaks into a union peak set and generated a peak-accessibility matrix using addPeakMatrix.

We identified cell-type marker peaks from the peak matrix using ArchR getMarkerFeatures with a bias-corrected Wilcoxon test, retaining peaks with FDR ≤0.01 and log2FC ≥1. For each species, we generated cell-type pseudobulk accessibility profiles using ArchR getGroupSE, calculating mean accessibility per cell. We CPM-normalized and log2-transformed the resulting profiles and restricted the analysis to the 25,000 most variable peaks. We performed PCA using cell types as samples.

We annotated transcription-factor motifs using the JASPAR2024 CORE non-redundant vertebrate position-frequency-matrix collection, converted the motifs to a PWMatrixList and added them to the ArchR projects using addMotifAnnotations (118). We assessed motif enrichment in marker peaks and differentially accessible peaks using peakAnnoEnrichment, retaining enriched motifs with FDR ≤0.1 and log2FC ≥0.5, and visualized the results as heatmaps.

For each species, we derived a cell-type transcription-factor motif-enrichment matrix from the ArchR motif-enrichment analysis. Because the same transcription-factor motif database was used for all species, these matrices were combined for cross-species analysis. We calculated pairwise Pearson correlations between cell types across the four species and visualized the resulting relationships using hierarchical clustering and heatmaps. For analyses of transcription-factor activity patterns across cell types, we converted the merged motif-enrichment matrix to ranked scores. We retained transcription factors with activity scores >2 in at least three cell types and set scores <0.2 to zero. We visualized the resulting patterns across cell types using line plots.

### Gene regulatory network reconstruction with SCENIC+

SCENIC+ integrates chromatin accessibility and gene-expression information to infer enhancer-driven gene regulatory networks. The framework combines pycisTopic, which uses topic modeling to identify groups of co-accessible regulatory regions, with cisTarget, which identifies transcription-factor binding motifs enriched in candidate regulatory regions, to construct enhancer–TF–gene regulatory relationships (eRegulons) (38). We applied SCENIC+ independently to each of the four species with snATAC-seq and snRNA-seq data, treating the two modalities as pseudo-multiome data.

For each species, we generated the genome and annotation files required by SCENIC+. We indexed each genome assembly using SAMtools v1.18 and extracted chromosome sizes from the resulting .fai files (109). We generated strand-aware TSS BED files from the GTF annotations, using the 5′-most coordinate for genes on the positive strand and the 3′-most coordinate for genes on the negative strand, together with a gene-annotation table. These files provided the genome sequence, TSS and gene-coordinate references required by the pycisTopic and SCENIC+ pipelines and served as the sequence source for the cisTarget databases.

We generated candidate regulatory regions and chromatin-accessibility topic models independently for each species using pycisTopic v2.0a0 (119). We exported cell-type labels and cell barcodes from the corresponding ArchR projects and pseudobulked Cell Ranger ATAC fragment files by cell type using export_pseudobulk. We called pseudobulk peaks with MACS2 v2.2.9.1 using BEDPE input, --shift 73, --ext_size 146, --keep-dup all and -q 0.05, with an effective genome size equal to the number of non-ambiguous bases in the genome assembly. We merged peaks into a consensus region set using get_consensus_peaks, specifying a peak half-width of 250 bp and no blacklist. We quality-filtered barcodes against the consensus regions and TSS annotation using pycistopic qc, applying the automatic unique-fragment and TSS-enrichment thresholds, and generated cisTopic objects using create_cistopic_object_from_fragments. We identified and removed doublets using Scrublet (120) with an expected doublet rate of 0.1 and a call threshold of 0.22.

We fitted latent Dirichlet allocation topic models using Mallet v202108 across models containing 2–50 topics, with 500 iterations, α = 50 per topic, η = 0.1 and seed 555 (121). We selected the optimal number of topics by manual assessment of multiple model-selection metrics, including the Griffiths–Steyvers, Cao, Arun and Mimno metrics (122–125). We clustered cells using Leiden clustering with k = 10 at resolutions of 0.6, 1.2 and 3 (126) and generated UMAP and t-SNE embeddings. We binarized topic-associated regions using the Otsu method, retaining the top 3,000 regions per topic.

We identified differentially accessible regions (DARs) between cell types using imputed accessibility. Accessibility values were imputed using impute_accessibility with a scale factor of 10^6 and normalized using normalize_scores with a scale factor of 10^4. We identified highly variable features using minimum dispersion of 0.05, minimum mean of 0.0125 and maximum mean of 3, and identified cell-type DARs using find_diff_features, retaining regions with adjusted P ≤0.05 and log2FC ≥log2(1.5). Binarized topic-associated regions and cell-type DARs were exported as BED files and used as candidate regulatory-region sets for SCENIC+ (38).

Because precomputed cisTarget motif databases are available only for human, mouse and Drosophila, we generated species-specific databases for each poeciliid species using the Aerts laboratory create_cisTarget_databases pipeline (38). We extracted the consensus regulatory regions from each genome with 1-kb flanking background sequence and scored them against the Aerts laboratory clustered motif collection (v10nr_clust_public singletons) using Cluster-Buster (127). This generated region-by-motif ranking and motif-score databases. We generated species-specific motif-to-transcription-factor annotation tables by mapping transcription factors identified using AnimalTFDB v4.0 (128) to human transcription factors using 1:1 orthologs inferred with OrthoFinder. We combined direct and orthology-based motif annotations to generate the motif annotation resources used for SCENIC+ (38, 128).

### SCENIC+ eRegulon inference

We inferred eRegulons independently for each species using SCENIC+ v1.0a2 and its Snakemake workflow (38). We treated the snATAC-seq and snRNA-seq atlases as pseudo-multiome data and aggregated cells into metacells containing 10 cells matched by cell-type label. We performed motif enrichment on candidate regulatory-region sets using both cisTarget and differential enrichment of motifs (DEM). For cisTarget, we used a normalized enrichment score (NES) threshold of 2.5, an area-under-the-curve (AUC) threshold of 0.005 and a rank threshold of 0.05. For DEM, we used adjusted P ≤0.1 and log2FC ≥0.5, with up to 500 background regions and promoter balancing. We used direct and orthology-based extended motif annotations, with a motif-similarity FDR threshold of 0.001. We relaxed several thresholds relative to the default SCENIC+ settings to maximize comparability across the four poeciliid species, with stringency imposed during the cross-species analyses. This procedure generated direct eRegulons, supported by experimentally characterized transcription-factor motifs, and extended eRegulons incorporating orthology-based motif assignments. For each species, we obtained gene-and region-based eRegulon AUC scores for each cell type. We calculated eRegulon specificity scores and visualized the ten highest-scoring activator eRegulons (+/+) for each cell type as heatmaps.

We constructed consensus gene-regulatory networks from the SCENIC+ eRegulon output, retaining direct activator (+/+) and repressor (−/−) eRegulons. Within each species, we considered a transcription factor (TF) active in a cell type when its binarized median gene-based AUC score was 1 and its pseudobulk expression exceeded 0.5. We retained TF–target interactions among the top 50% of triplet-confidence ranks and restricted transcription factors to those active in no more than five cell types to exclude broadly active, non-specific regulons. We mapped retained regulons into a common P. turneri orthology space using 1:1 OrthoFinder orthologs. For each cell type, we calculated a consensus activity score for each transcription factor as the number of species in which the corresponding regulon was active. We visualized the eight transcription factors with the highest consensus activity per cell type as regulatory networks. Interaction color indicated regulatory direction, with activation and repression represented separately; interaction width was scaled according to interaction confidence (100 − mean triplet-rank percentile), and node color indicated the number of species in which the transcription factor was active.

### Conservation of regulatory control of differentially expressed genes

We asked whether the regulatory interactions controlling genes up-regulated in the late follicle cells of each placental species were already present in its non-placental sister species. We tested this for *P. turneri* against *P. gracilis*, and for *P. bifurca* against *P. picta*. Using the set of DEGs, we took genes assigned to the placental lineage with log2 fold change > 1 on that lineage and adjusted P < 0.05 for the placental–sister comparison.

We built the placental regulatory network from the late follicle cell-specific direct activator (+/+) eRegulons. We kept only eRegulons active in the late follicle cell, meaning a binarized median gene-based AUC score of 1. We also kept only TF–target interactions within the top 80% of triplet-confidence ranks, and target genes with pseudobulk expression ≥ 0.1 in the late follicle cell. We then kept the interactions whose target was a placental-lineage DEG. We translated each transcription factor and its target into the sister species using 1:1 OrthoFinder orthologs. We classed an interaction as untestable if its transcription factor lacked a 1:1 ortholog, and excluded it. We compared each testable interaction with the sister species’ full SCENIC+ TF–gene network across all cell types, and with its late follicle cell +/+ eRegulons. An interaction was “pre-existing” if it appeared among the sister late follicle cell eRegulons or if it appeared elsewhere in the sister network (in another cell type). Otherwise, we called it a “novel interaction”.

### Positive selection analysis

We translated the coding sequences of 14,204 1:1 orthologous genes into peptide sequences using AGAT v1.0 and filtered the resulting sequences for poorly aligned and ambiguous regions using PreQual v1.02 (129). We aligned the filtered peptide sequences using PRANK (130), a phylogeny-aware alignment method, and converted the resulting protein alignments to codon-based nucleotide alignments using PAL2NAL v14.1 (131). We tested for signatures of positive selection at individual codon sites using the site models implemented in codeml within PAML v4.10.7 (132). For each orthogroup, we compared the nearly neutral M1a model, which allows sites to evolve under purifying (dN/dS < 1) or neutral (dN/dS = 1) selection, with the alternative M2a model, which additionally allows a proportion of sites to evolve under positive selection (dN/dS > 1). We used a fixed species-tree topology and applied the same species tree to all orthogroups. We compared model fit using a likelihood-ratio test, calculated as twice the difference in log-likelihood between the M2a and M1a models, and evaluated the statistic against a χ² distribution with 2 degrees of freedom. We considered orthogroups with a significant likelihood-ratio test after Benjamini–Hochberg false-discovery-rate correction (FDR-adjusted P < 0.05) as candidates for positive selection.

### Parallel substitution analysis

We tested for convergent evolution at the codon level using CSUBST v1.4.20 (45), which identifies the co-occurrence of the same or similar nonsynonymous substitutions on independent branches of a phylogeny as evidence of convergent or parallel amino-acid evolution. We used the codon alignments generated with PAL2NAL as input. For each orthogroup, we performed ancestral sequence reconstruction and branch-wise substitution mapping using IQ-TREE under the GY+F+G4 codon substitution model (82). We used a fixed, rooted species-tree topology for all orthogroups. We defined *P. turneri*, *P. prolifica*, *H. formosa* and *P. bifurca* as foreground lineages (placental lineages) and *P. gracilis*, *P. infans*, *B. rhabdophora* and *P. picta* as background lineages (non-placental lineages).

We identified candidate genes using two sequential filtering criteria. First, we retained only orthogroups with a median gene-level combinatorial substitution rate ratio (omega_C), calculated across all pairwise comparisons among foreground species, of ≥ 2.0. Second, among these orthogroups, we identified individual nonsynonymous substitutions that occurred on at least two foreground branches and were absent from all background branches. We excluded genes with fewer than two such foreground-exclusive nonsynonymous substitutions. Orthogroups passing both criteria were considered candidate genes exhibiting convergent amino acid substitution among the foreground lineages. As a null expectation, we repeated the analysis after reversing the foreground and background lineage assignments.

### Gene-age inference

We estimated the evolutionary age of protein-coding genes independently for each genome using GageTracker (44). GageTracker estimates gene age by identifying conserved genomic regions across a phylogenetically ordered set of outgroup genomes using micro-and macro-synteny. For each species, we supplied the genome assembly, gene annotation and a set of outgroup genomes in a species-specific control file. We arranged the outgroup species into phylogenetic branches from older to younger evolutionary nodes and used these branch assignments to determine the oldest evolutionary interval in which each gene could be detected. We ran GageTracker with 10 parallel processes and masked repetitive regions in the outgroup genomes using the -mos option. We additionally used the -m option to infer the putative mechanism of origin for young genes based on protein-sequence similarity. We performed gene-age estimation independently for each species using the same general procedure, with the outgroup composition and phylogenetic branch assignments specified for each target genome. GageTracker reports the inferred age of each gene together with a confidence classification based on the underlying genome alignments. We retained the gene-age assignments generated from the final GageTracker output for downstream analyses of the evolutionary age and origin of genes across species.

## Acknowledgments

We thank R. Arguello, C. Collart, R. Enriquez-Gasca, C. Jones and G. Taylor for comments on the manuscript; D. Snell, H. Slawinski, and the Genomics facility at the Francis Crick Institute for sequencing reads and library preparations; M. Millington and her team at the Aquatics facility for maintaining and breeding poeciliids.

## Author contributions

D.S., S.K. and M.C.-M. conceived the study. D.S. and S.K. sampled, generated and analysed all the data. M.F. developed pipelines for cross-species comparisons. D.R. and L.L. maintained and provided fish. D.S., S.K. and M.C.-M. wrote the manuscript with contributions from all authors. A coin toss decided the order of the two first authors.

## Data, code, and materials availability

The raw and processed data generated in this study for the genome assemblies were deposited in the European Nucleotide Archive (ENA) with the accession code PRJEB115767. The snRNA-seq and snATAC-seq raw and processed data were deposited in ArrayExpress with the accession codes E-MTAB-17757 and E-MTAB-17776, respectively. The code used to analyze the data, the list of1:1 orthologs, the list of genes under positive selection and the list of genes with parallel substitution are available at https://github.com/cardosomoreiralab/fish-placentas-2026. Processed data can be interactively explored at https://shiny.cardosomoreiralab.org/fishplacentas/.

---

**Supplementary figure 1.**
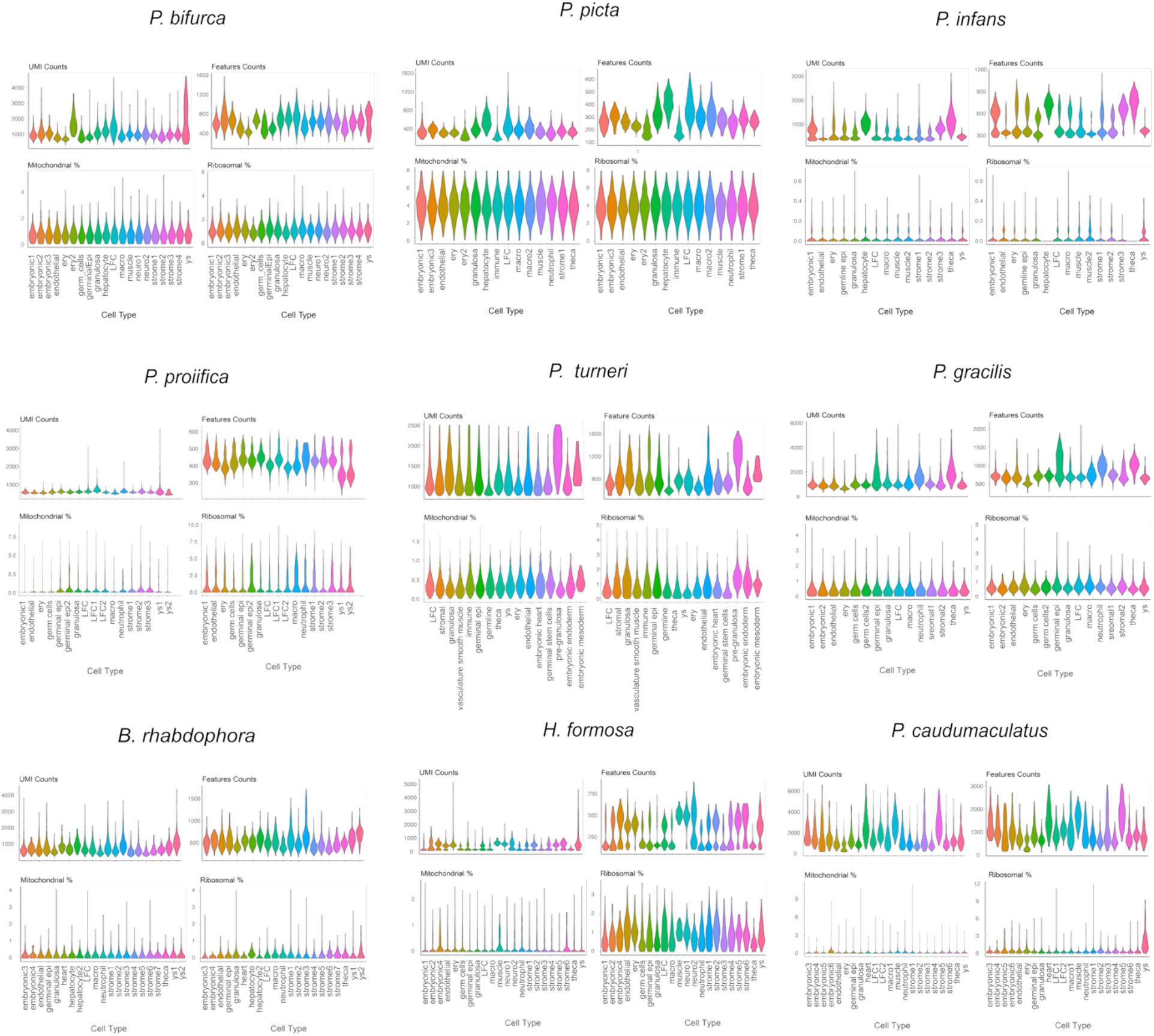
Characteristics of the snRNA-seq atlases across species.

**Supplementary figure 2.**
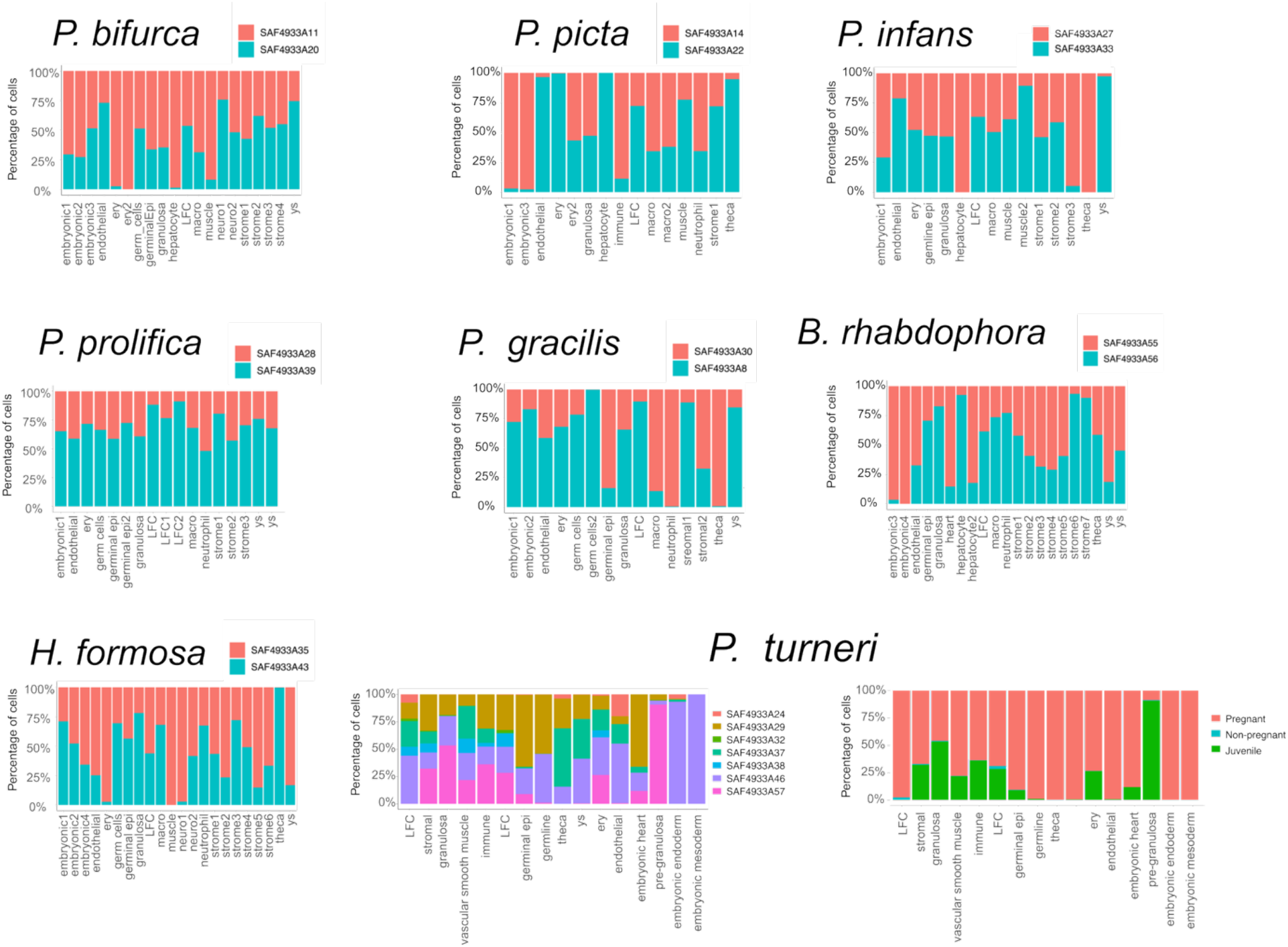
Contribution of each snRNA-seq library to the species atlas.

**Supplementary figure 3.**
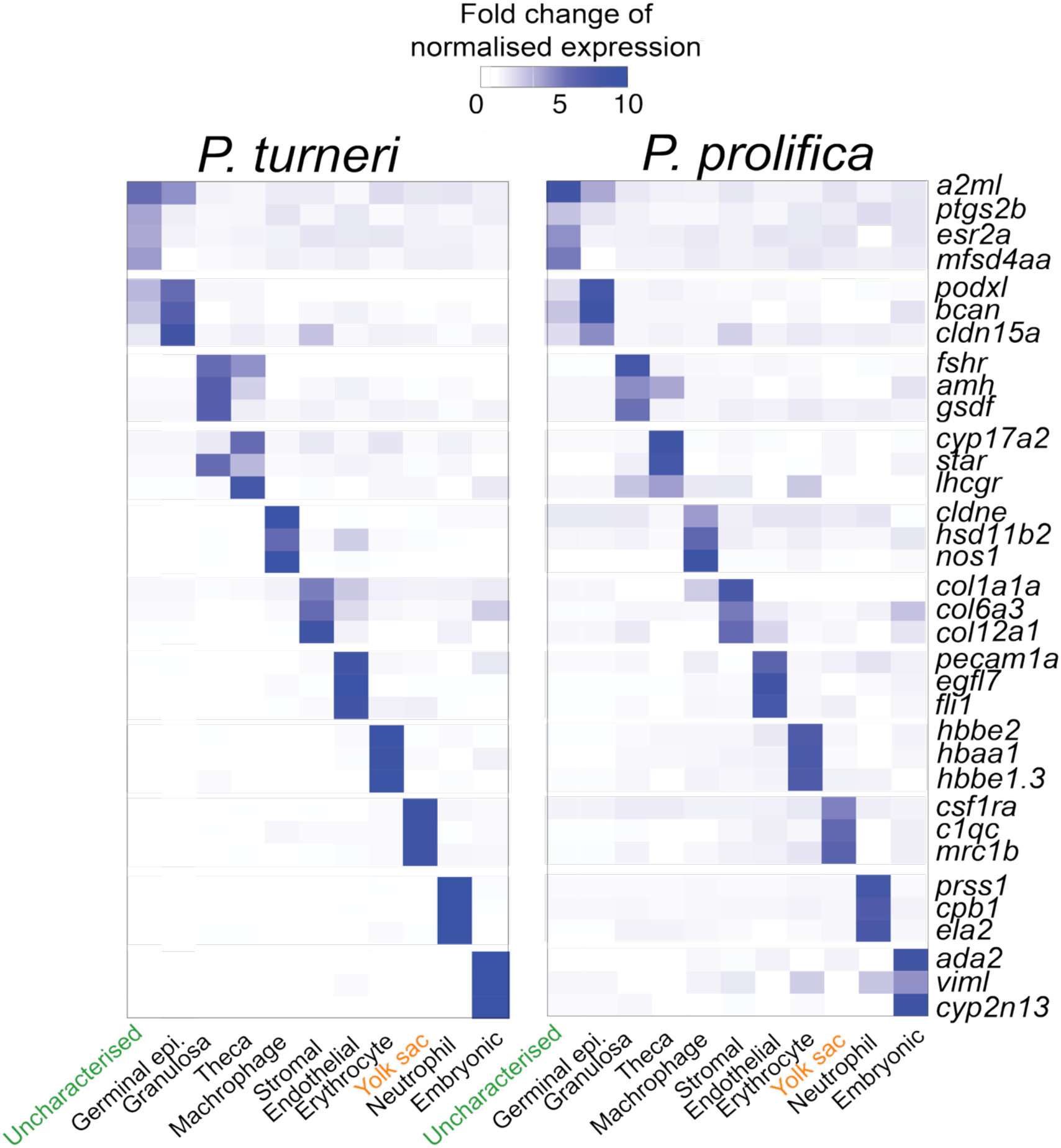
Cell type-specific gene markers for ovarian cell types in *P. turneri* and *P. prolifica*.

**Figure Supplementary 4.**
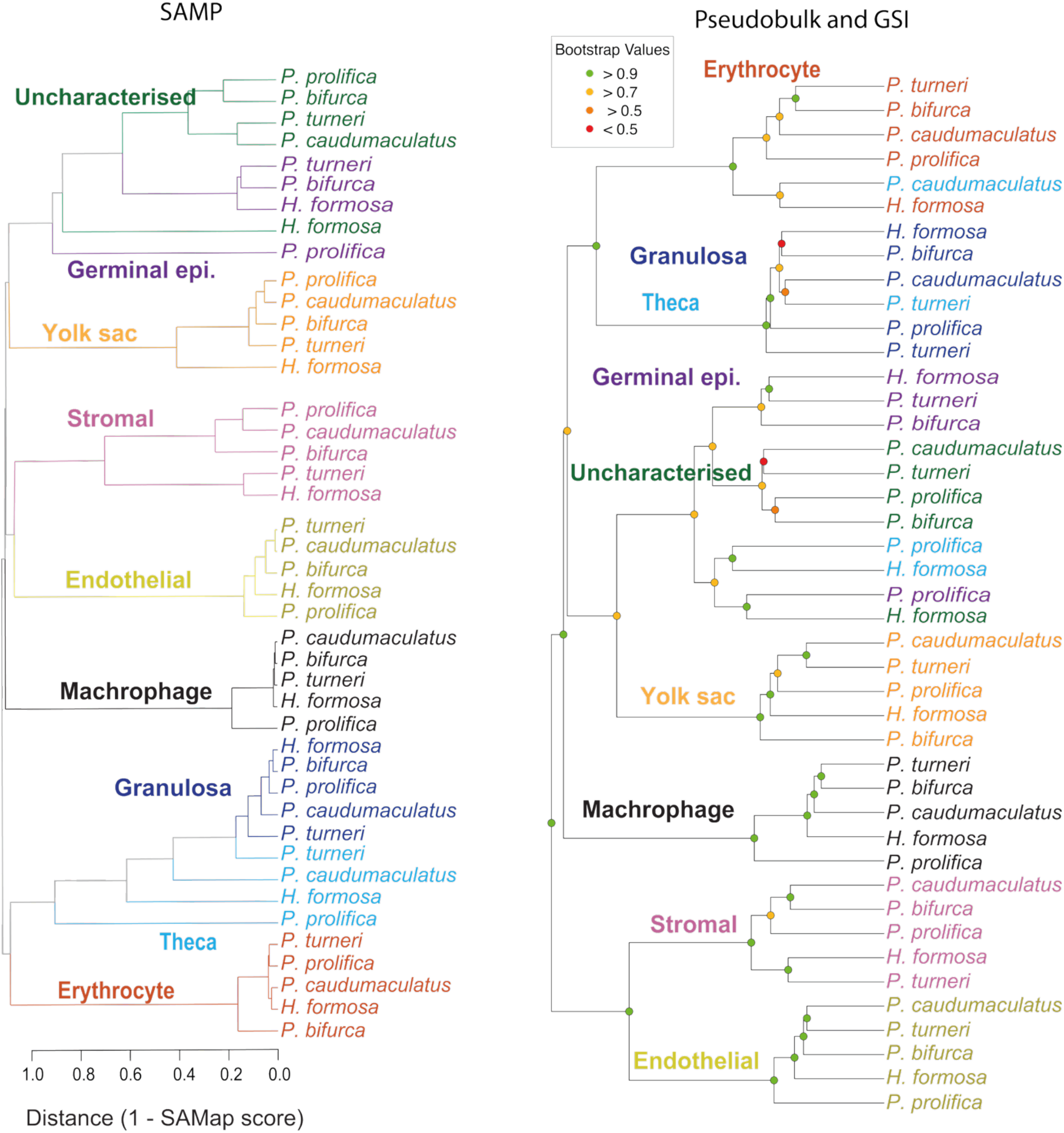
Dendrograms based on distance scores (1 - SAMap score) (left) and based on pseudobulk and GSI (right) for ovarian cells from the five placental species.

**Supplementary figure 5.**
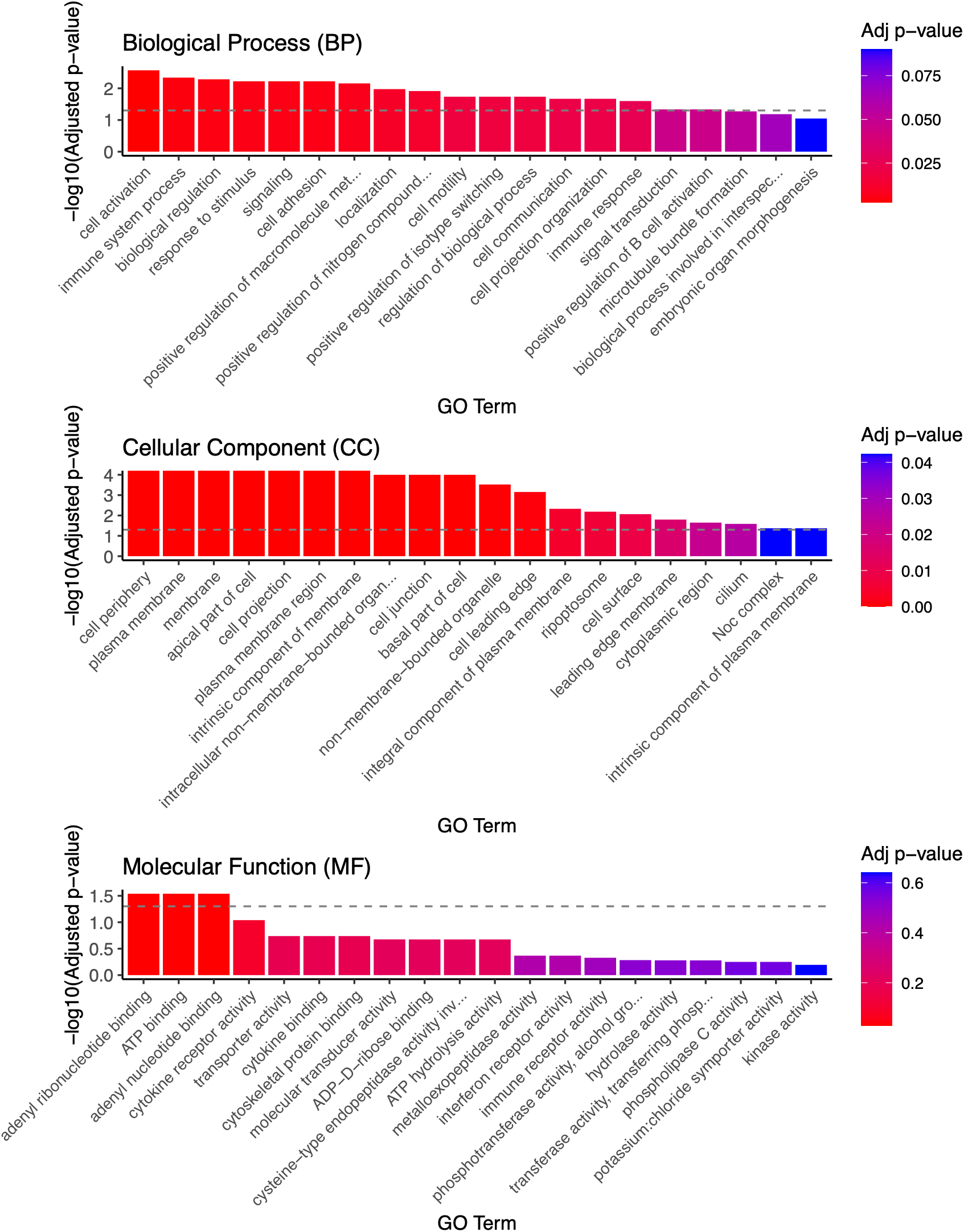
GO enrichments of the genes under positive selection across poeciliids. Dotted line represents P < 0.05.

**Supplementary figure 6.**
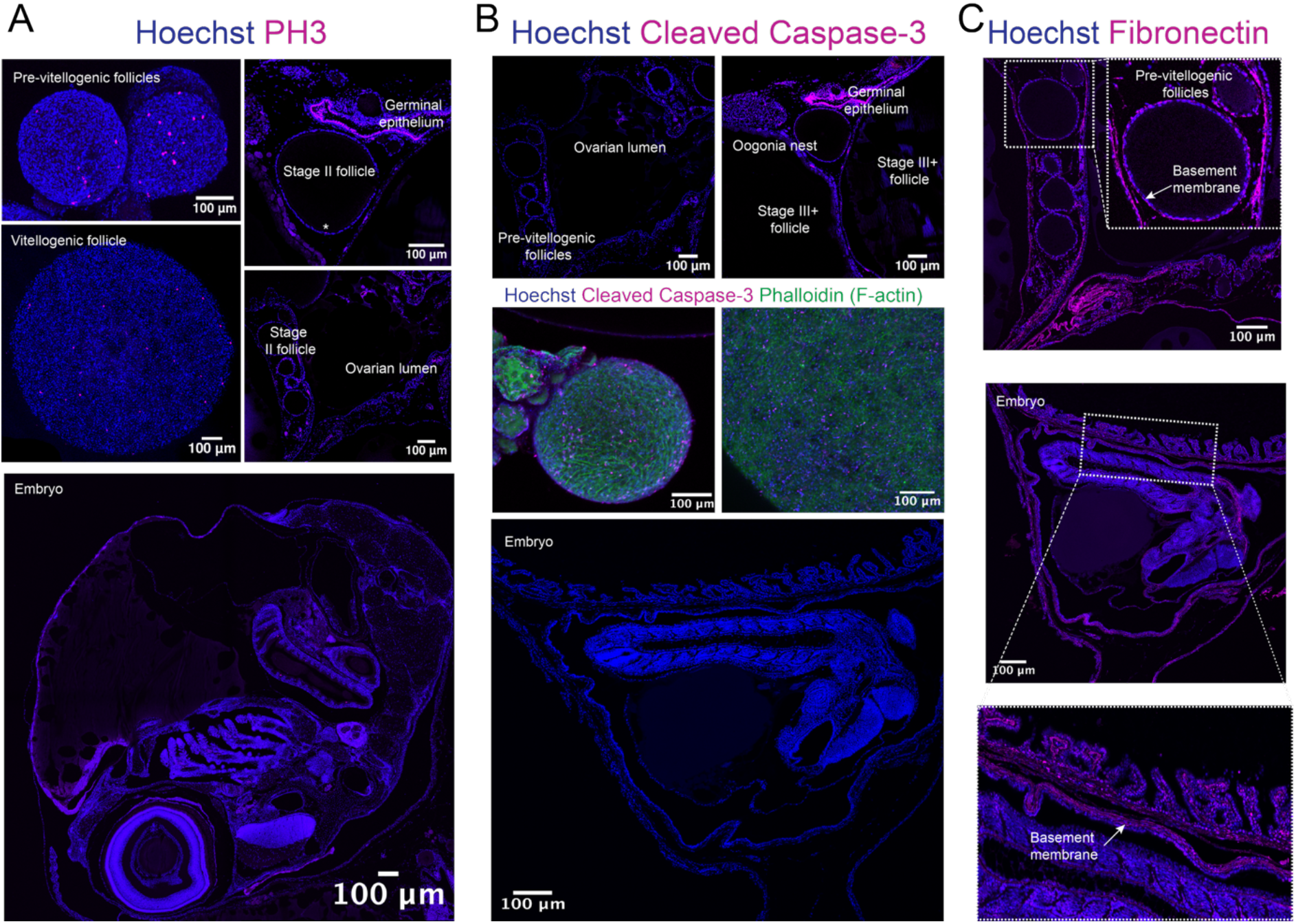
Immunofluorescence for **(A)** proliferation (PH3), **(B)** apoptosis (cleaved Caspase-3) and **(C)** basement membrane (Fibronectin).

**Supplementary figure 7.**
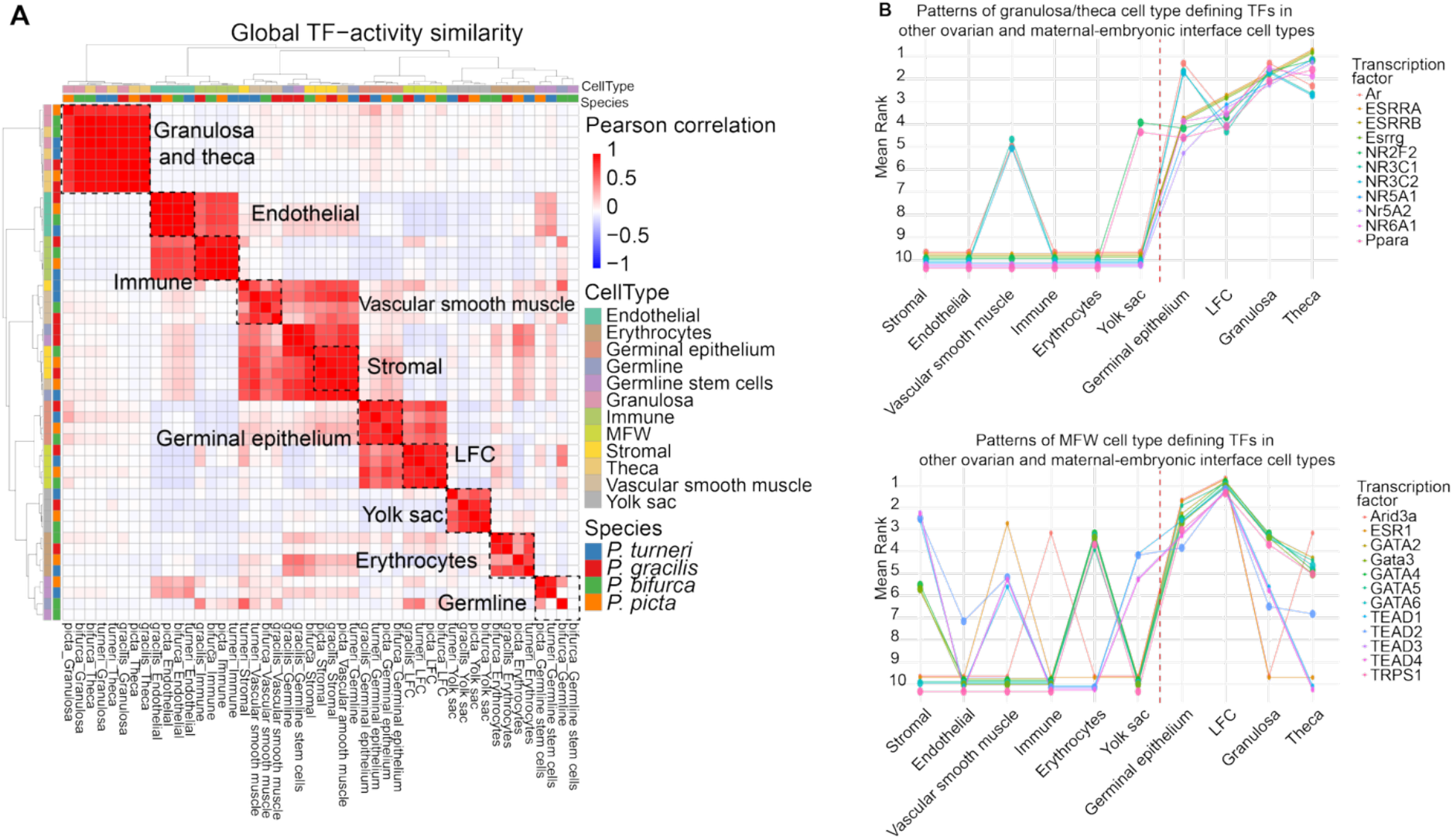
**(A)** Heatmaps showing hierarchical clustering (Ward’s D2) of pairwise Pearson correlations for global TF activity, inferred from motif enrichment in cell-type marker peaks. **(B)** Line plot of TF activity rank across ovarian and maternal–embryonic interface cell types for TFs most active in granulosa/theca or late follicle/germinal epithelium cells. TFs were included if active (activity > 2) in at least three cell types, with initial ranking conditions met in at least two species. Dot size indicates the number of species supporting each ranking.

**Supplementary figure 8.**
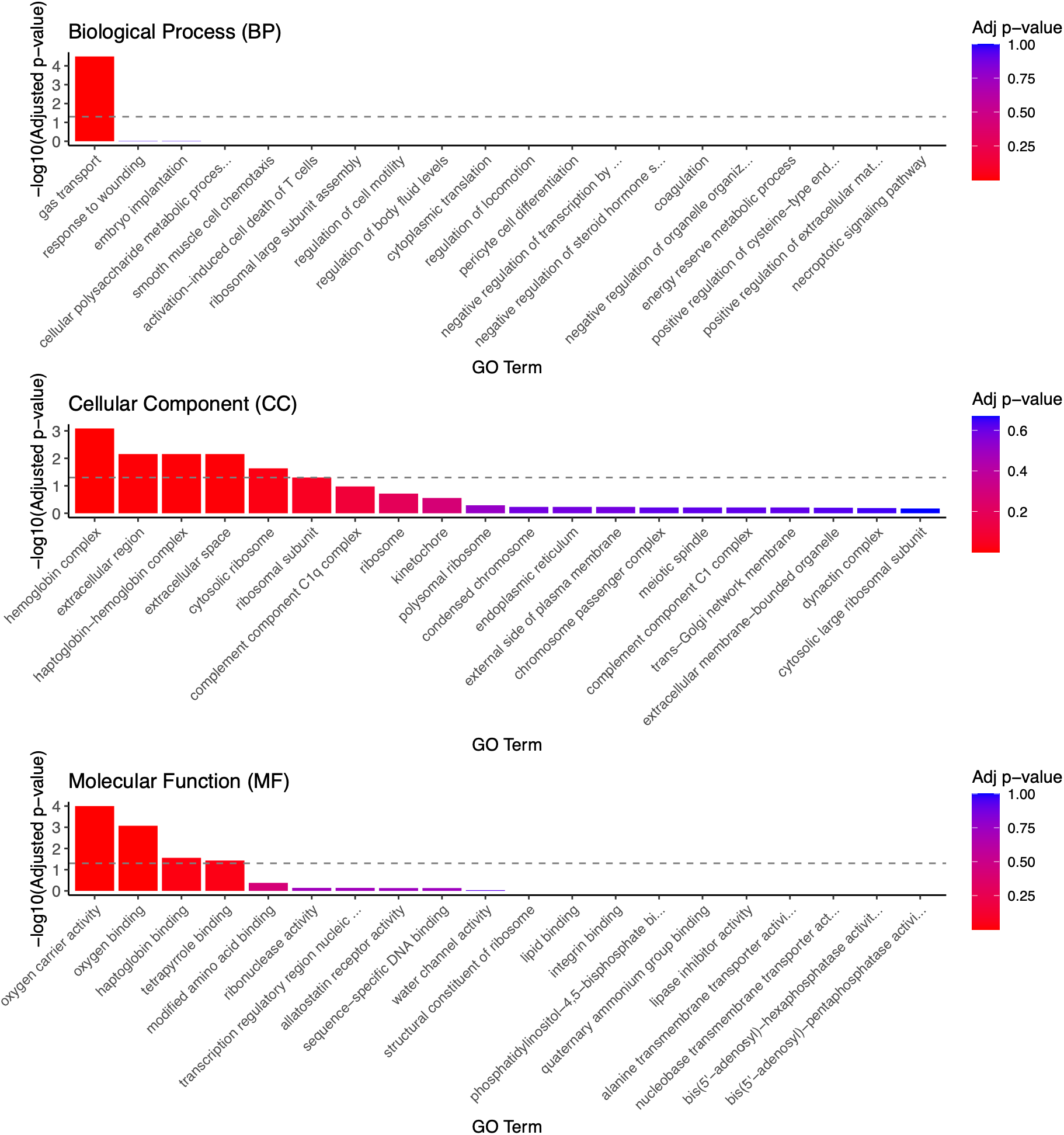
GO enrichment of differentially expressed genes in the late follicle cell type of *B. rhabdophora.* Dotted line represents P < 0.05.

**Supplementary figure 9.**
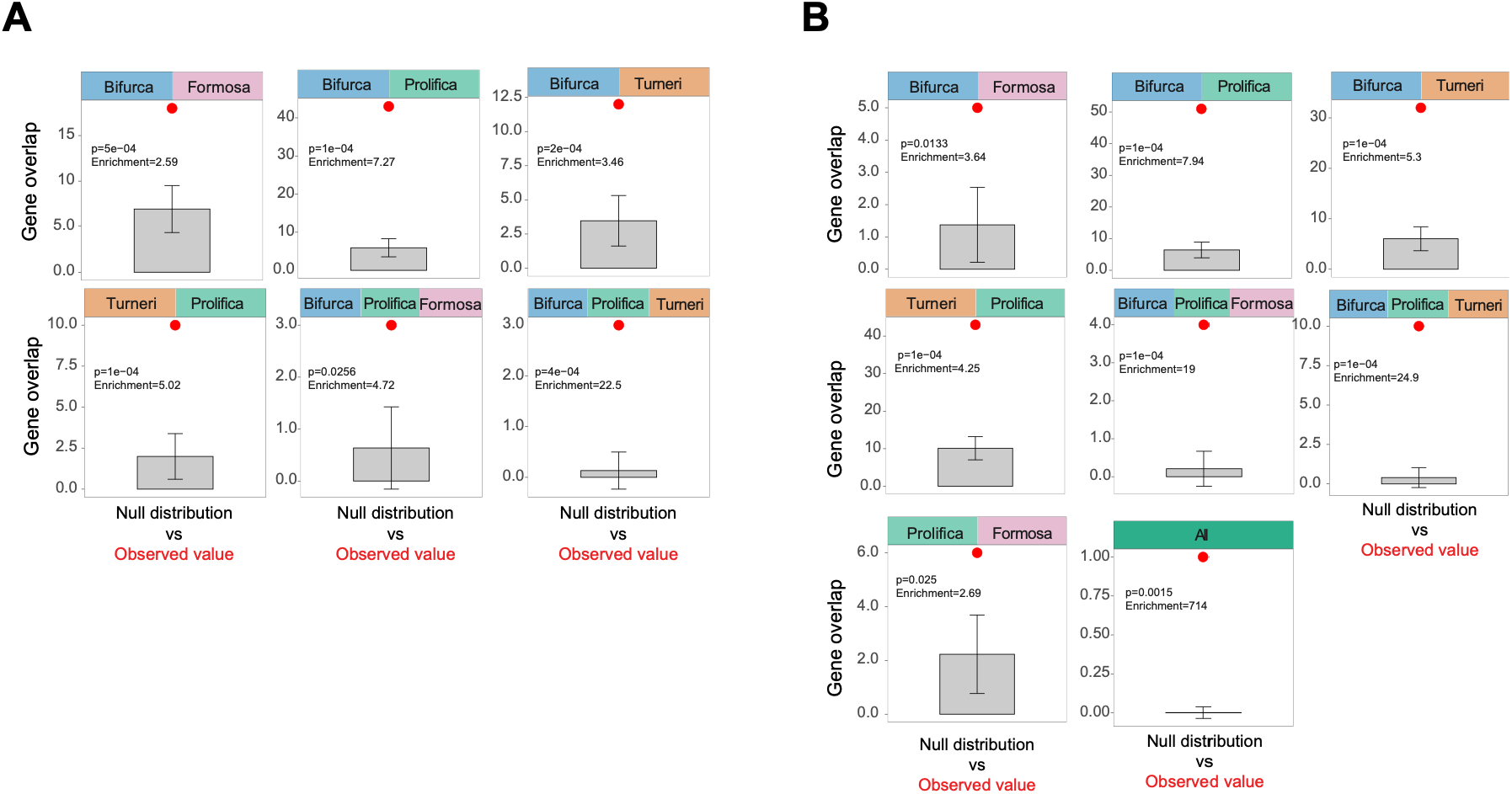
Pairwise overlap of differentially expressed genes in the late follicle cell type across placental species. **(A)** Pairwise overlap of upregulated genes. **(B)** Pairwise overlap of downregulated genes. Observed overlaps were compared with null distributions generated by permutation testing to assess whether shared gene counts exceeded random expectations. Statistical significance of pairwise overlaps was assessed using one-sided permutation tests with 10,000 permutations. Empirical P values were calculated as the proportion of permutations yielding an overlap equal to or greater than the observed overlap. Enrichment was calculated as the ratio of the observed overlap to the mean overlap expected under the permutation-based null distribution.

**Supplementary figure 10.**
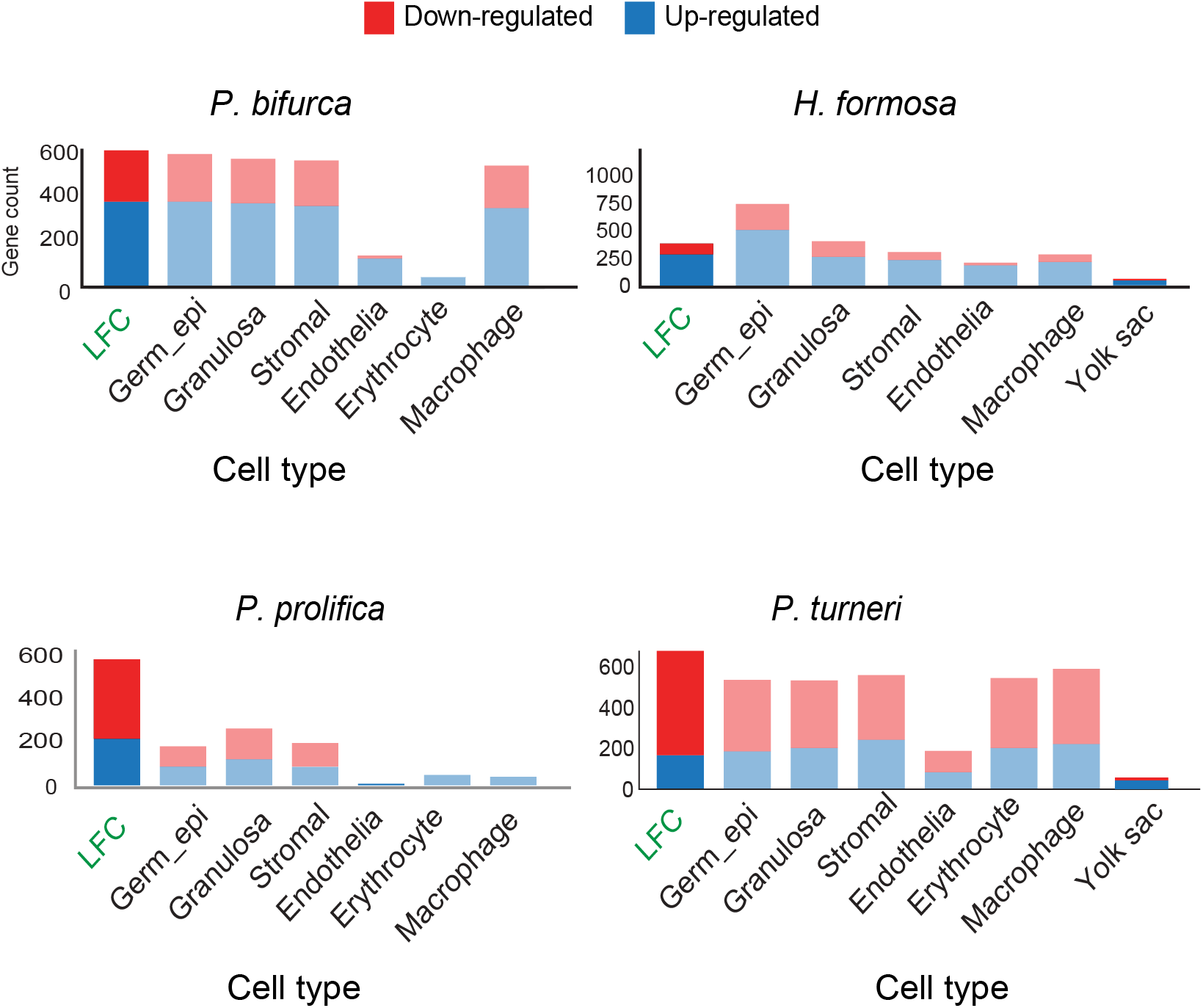
Number of differentially expressed genes in placental species across cell types.

**Supplementary figure 11.**
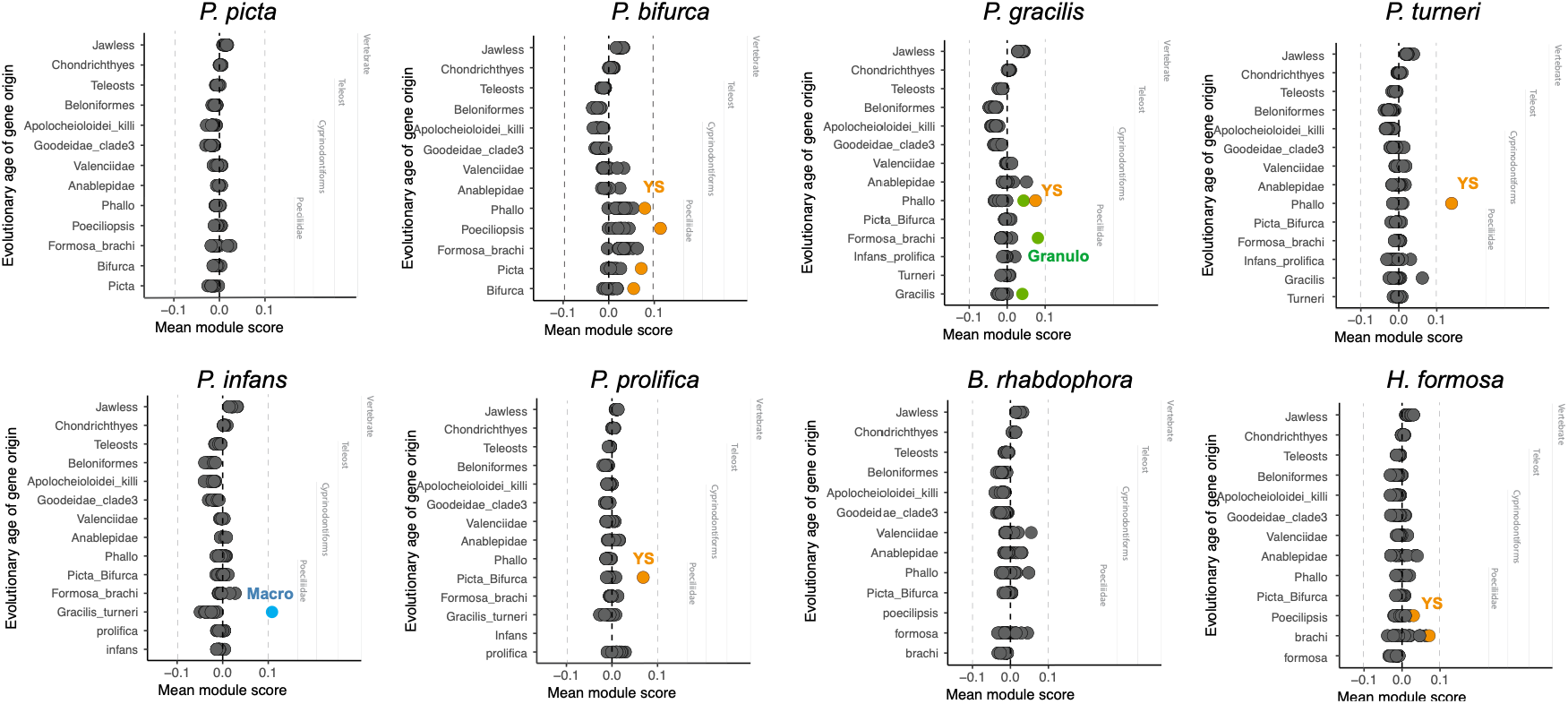
Enrichment scores for genes with different evolutionary ages across cell types in all four placental-non-placental pairs. *P. picta, P. infans, P. gracilis* and *B. rhabdophora* correspond to the non-placental pairs for the placental species *P. bifurca, P. prolifica. P turneri* and *H. formosa*, respectively.

**Supplementary table 1.**
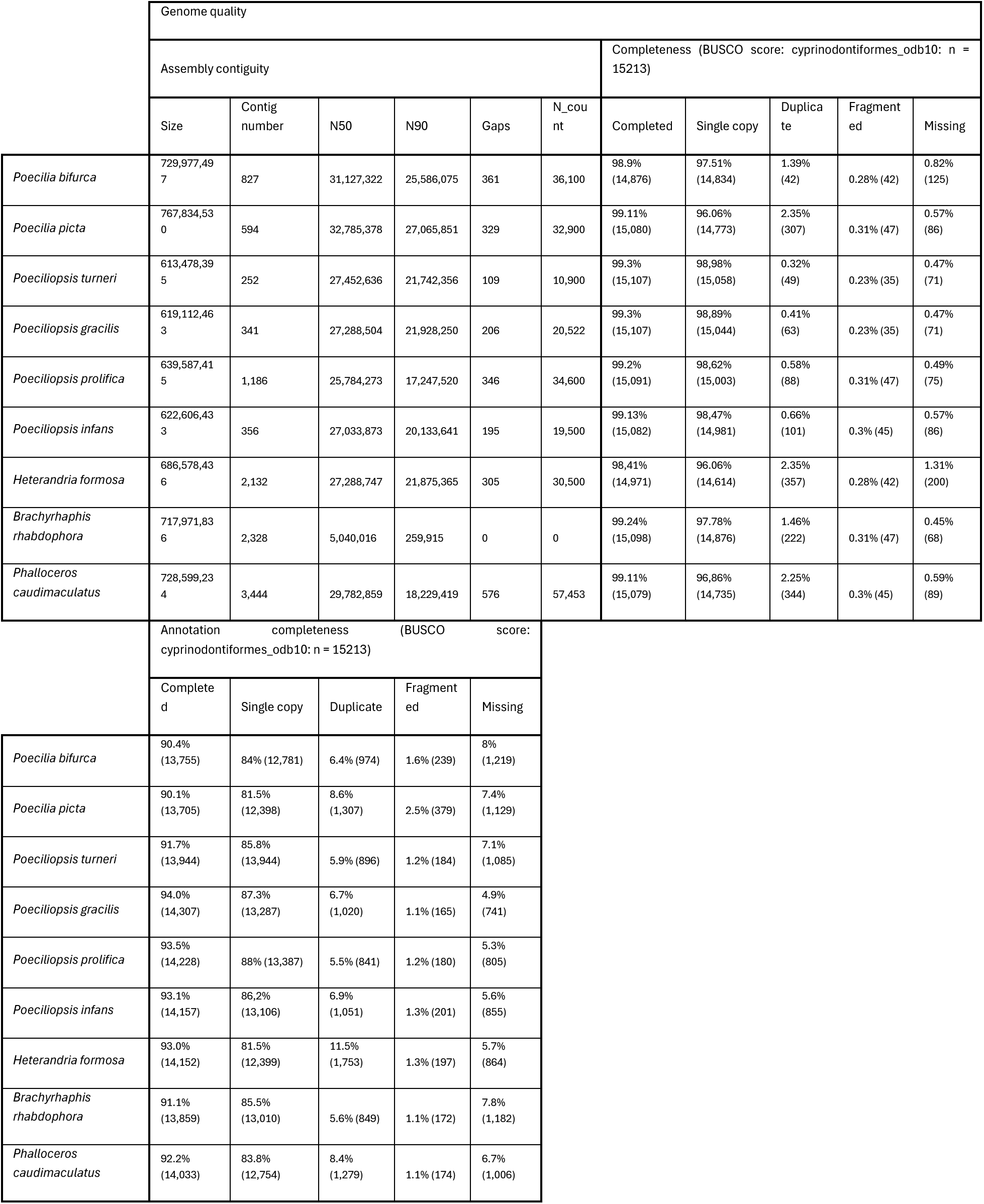
Summary of genome assembly quality and annotation completeness across the nine poeciliid species included in this study.

|  | Genome quality |  |  |  |  |  |  |  |  |  |
|---|---|---|---|---|---|---|---|---|---|---|
|  | Assembly contiguity |  |  |  |  |  | Completeness (BUSCO score: cyprinodontiformes_odb10: n = 15213) |  |  |  |
|  | Size | Contig number | N50 | N90 | Gaps | N_count | Completed | Single copy | Duplicate | Fragmented | Missing |
| *Poecilia bifurca* | 729,977,497 | 827 | 31,127,322 | 25,586,075 | 361 | 36,100 | 98.9% (14,876) | 97.51% (14,834) | 1.39% (42) | 0.28% (42) | 0.82% (125) |
| *Poecilia picta* | 767,834,530 | 594 | 32,785,378 | 27,065,851 | 329 | 32,900 | 99.11% (15,080) | 96.06% (14,773) | 2.35% (307) | 0.31% (47) | 0.57% (86) |
| *Poeciliopsis turneri* | 613,478,395 | 252 | 27,452,636 | 21,742,356 | 109 | 10,900 | 99.3% (15,107) | 98,98% (15,058) | 0.32% (49) | 0.23% (35) | 0.47% (71) |
| *Poeciliopsis gracilis* | 619,112,463 | 341 | 27,288,504 | 21,928,250 | 206 | 20,522 | 99.3% (15,107) | 98,89% (15,044) | 0.41% (63) | 0.23% (35) | 0.47% (71) |
| *Poeciliopsis prolifica* | 639,587,415 | 1,186 | 25,784,273 | 17,247,520 | 346 | 34,600 | 99.2% (15,091) | 98,62% (15,003) | 0.58% (88) | 0.31% (47) | 0.49% (75) |
| *Poeciliopsis infans* | 622,606,433 | 356 | 27,033,873 | 20,133,641 | 195 | 19,500 | 99.13% (15,082) | 98,47% (14,981) | 0.66% (101) | 0.3% (45) | 0.57% (86) |
| *Heterandria formosa* | 686,578,436 | 2,132 | 27,288,747 | 21,875,365 | 305 | 30,500 | 98,41% (14,971) | 96.06% (14,614) | 2.35% (357) | 0.28% (42) | 1.31% (200) |
| *Brachyrhaphis rhabdophora* | 717,971,836 | 2,328 | 5,040,016 | 259,915 | 0 | 0 | 99.24% (15,098) | 97.78% (14,876) | 1.46% (222) | 0.31% (47) | 0.45% (68) |
| *Phalloceros caudimaculatus* | 728,599,234 | 3,444 | 29,782,859 | 18,229,419 | 576 | 57,453 | 99.11% (15,079) | 96,86% (14,735) | 2.25% (344) | 0.3% (45) | 0.59% (89) |
|  | Annotation completeness (BUSCO score: cyprinodontiformes_odb10: n = 15213) |  |  |  |  |  |  |  |  |  |
|  | Completed | Single copy | Duplicate | Fragmented | Missing |  |  |  |  |  |
| *Poecilia bifurca* | 90.4% (13,755) | 84% (12,781) | 6.4% (974) | 1.6% (239) | 8% (1,219) |  |  |  |  |  |
| *Poecilia picta* | 90.1% (13,705) | 81.5% (12,398) | 8.6% (1,307) | 2.5% (379) | 7.4% (1,129) |  |  |  |  |  |
| *Poeciliopsis turneri* | 91.7% (13,944) | 85.8% (13,944) | 5.9% (896) | 1.2% (184) | 7.1% (1,085) |  |  |  |  |  |
| *Poeciliopsis gracilis* | 94.0% (14,307) | 87.3% (13,287) | 6.7% (1,020) | 1.1% (165) | 4.9% (741) |  |  |  |  |  |
| *Poeciliopsis prolifica* | 93.5% (14,228) | 88% (13,387) | 5.5% (841) | 1.2% (180) | 5.3% (805) |  |  |  |  |  |
| *Poeciliopsis infans* | 93.1% (14,157) | 86,2% (13,106) | 6.9% (1,051) | 1.3% (201) | 5.6% (855) |  |  |  |  |  |
| *Heterandria formosa* | 93.0% (14,152) | 81.5% (12,399) | 11.5% (1,753) | 1.3% (197) | 5.7% (864) |  |  |  |  |  |
| *Brachyrhaphis rhabdophora* | 91.1% (13,859) | 85.5% (13,010) | 5.6% (849) | 1.1% (172) | 7.8% (1,182) |  |  |  |  |  |
| *Phalloceros caudimaculatus* | 92.2% (14,033) | 83.8% (12,754) | 8.4% (1,279) | 1.1% (174) | 6.7% (1,006) |  |  |  |  |  |

**Supplementary table 2.**
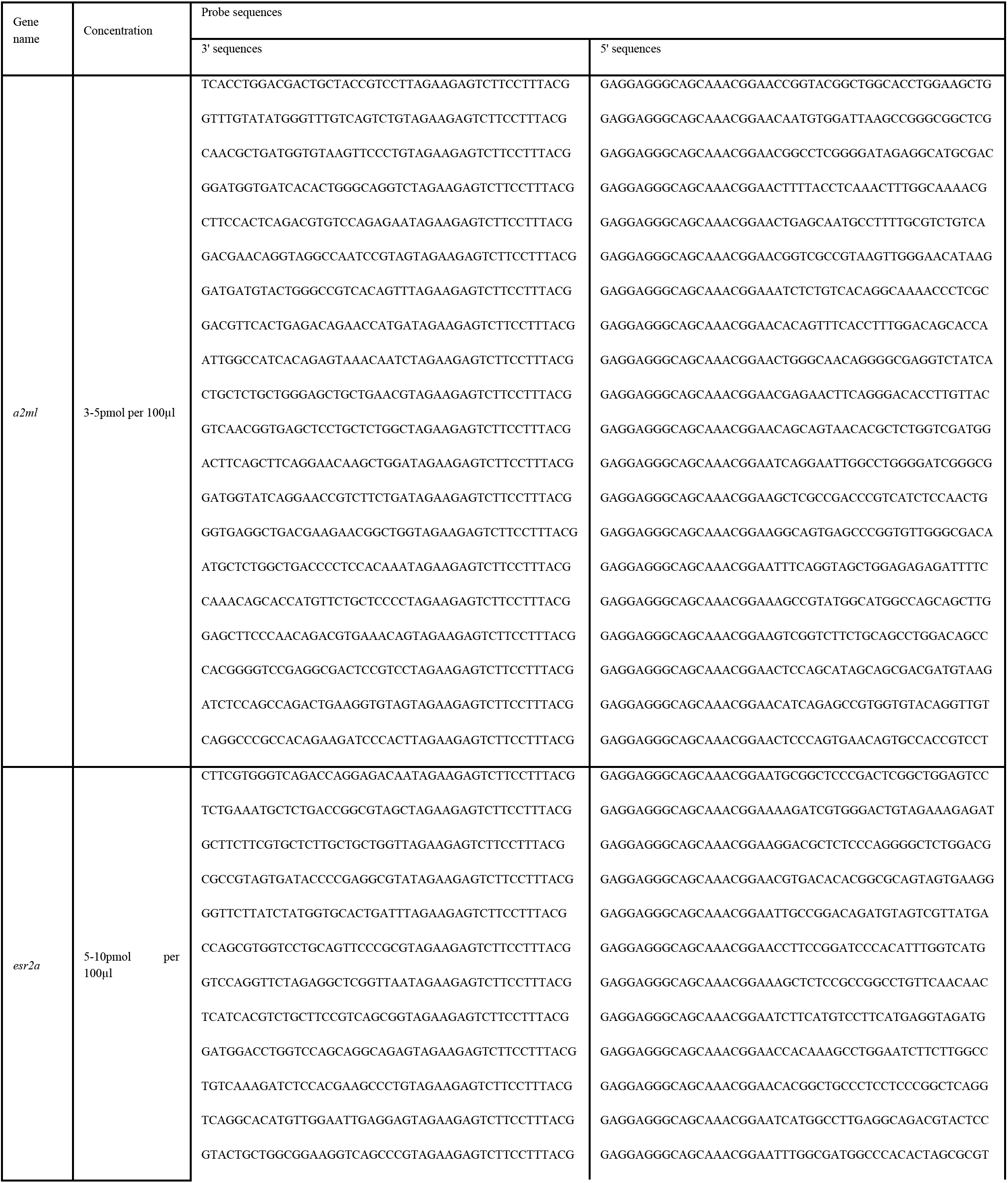
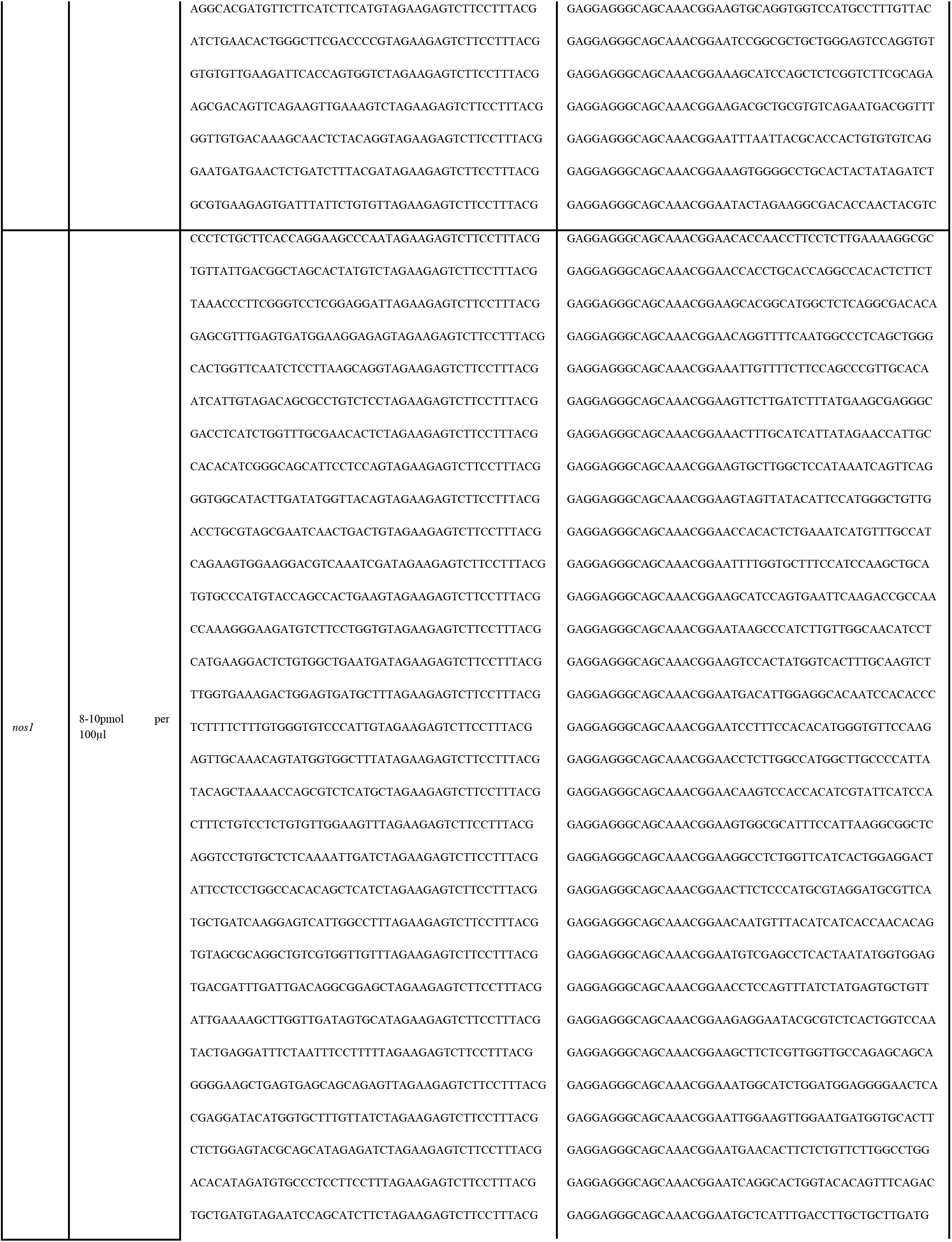
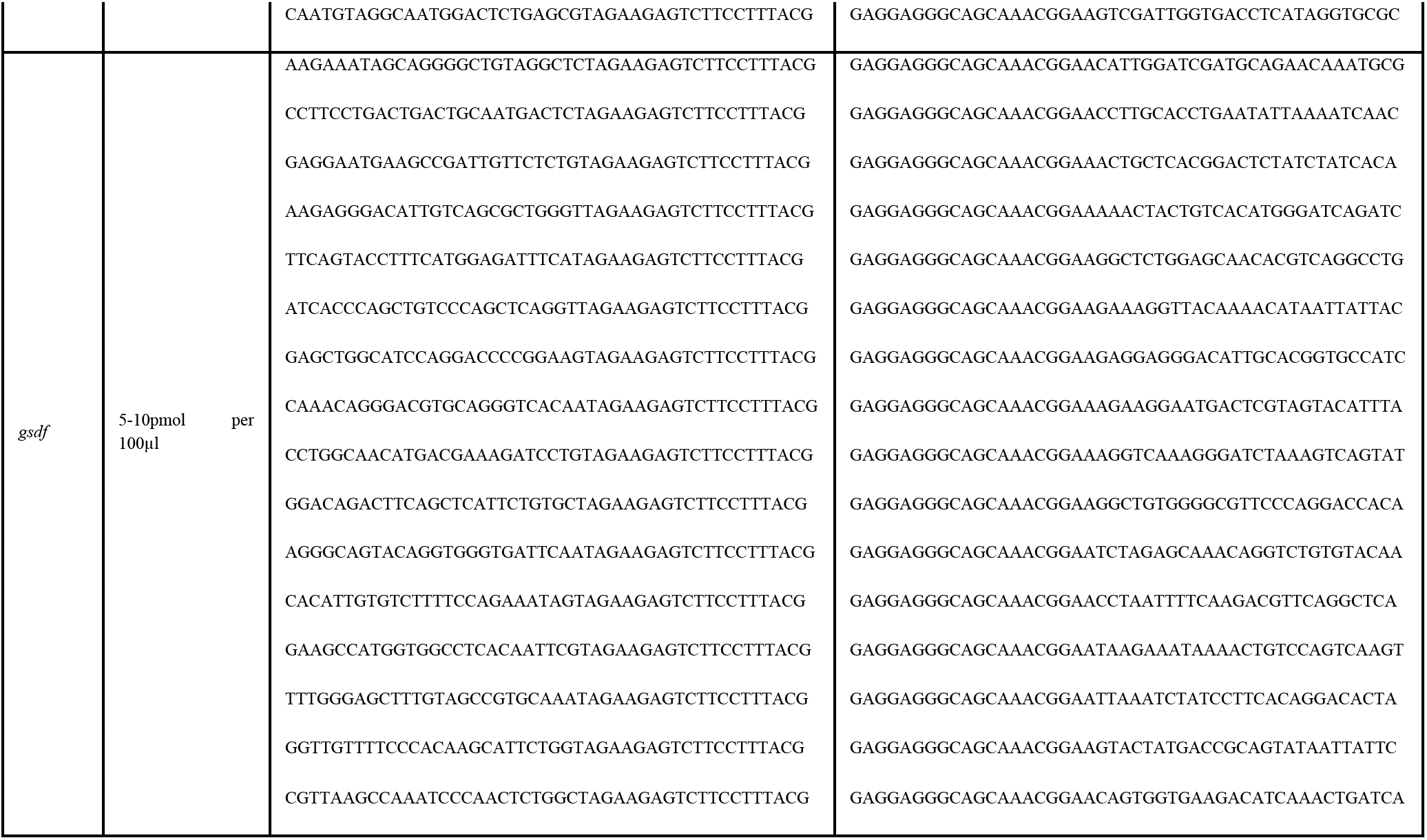
Hybridization chain reaction (HCR) probes used for gene expression analysis. The table lists the target gene (*a2ml*), probe concentration, and the 3′ and 5′ sequences of the oligonucleotide probes used for HCR.

**Supplementary table 3.**
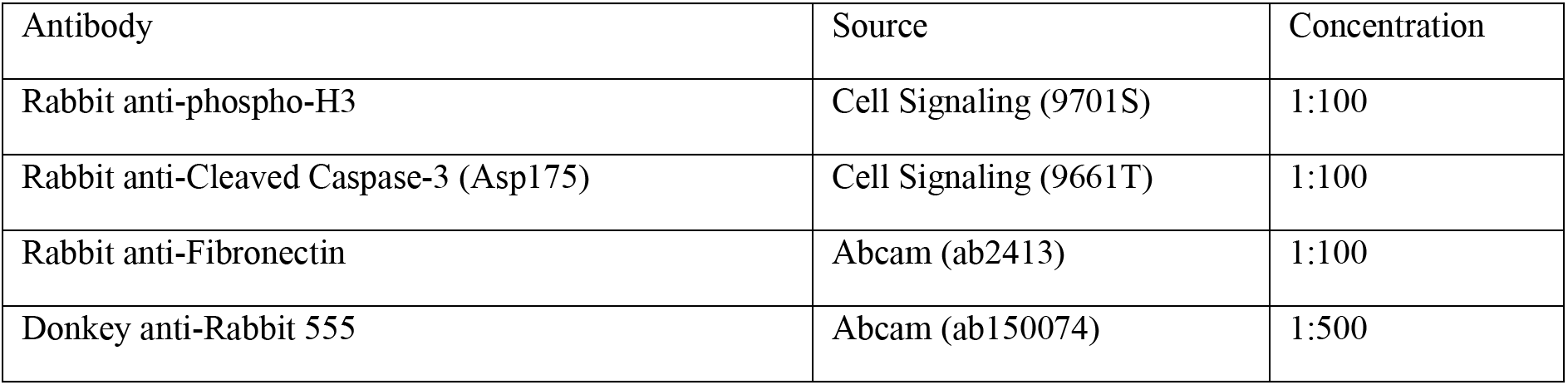
Information on antibodies used for immunofluorescence staining, including antibody specificity, source (manufacturer and catalog number), and working concentration used in the experiments.

